# Inferring the ecological drivers of arboviral outbreaks

**DOI:** 10.1101/632133

**Authors:** Warren Tennant, Trevelyan McKinley, Mario Recker

**Affiliations:** The Zeeman Institute: SBIDER, University of Warwick, Coventry, United Kingdom; Centre for Mathematics and the Environment, University of Exeter, Penryn Campus, Penryn, United Kingdom

## Abstract

The emergence and wide-spread circulation of mosquito-transmitted viral diseases, such as dengue, Zika and Chikungunya, is a global public health concern. In the absence of effective vaccines, current control measures are mostly targeted against the mosquito vector and have so far only shown limited success. The reliance on mosquitoes for transmission also imposes strong ecological constraints that can introduce significant spatial and temporal variations in disease incidence. However, the way that epidemiological and ecological factors interact and determine population-level disease dynamics is only partially understood. Here we fit a spatially-explicit individual based model defined within a Bayesian framework to Zika incidence data from Feira de Santana, allowing us to more precisely quantify the relationships between socio-ecological factors and arboviral outbreaks. Our results further demonstrated that the virus was likely introduced into multiple spatially segregated locations at the start of the outbreak, highlighting the benefits that spatio-temporal incidence data would bring in making modelling approaches more realistic for public health planning.

## 1 Introduction

Climate has been shown to play a key role in arboviral transmission dynamics as annual oscillations in temperature, precipitation and humidity induce seasonal fluctuations in vector suitability and virus transmissibility (Caminade et al., 2017; Johansson et al., 2009; Li et al., 2019). Increased temperatures have been associated with faster virus replication rates and shorter extrinsic incubation periods for Zika (Tesla et al., 2018), Chikungunya (Mbaika et al., 2016), dengue (Mordecai et al., 2017; Xiao et al., 2014), and yellow fever (Johansson et al., 2010). Higher temperatures also increase survivorship of mosquitoes, including *Aedes* (Alto and Bettinardi, 2013; Alto and Juliano, 2001), *Anopholes* (Lyons et al., 2013) and *Culex* (Ciota et al., 2014) species, the three most common genera of mosquito-borne disease vectors (Erlanger et al., 2009; Gubler, 2009; Jupp et al., 2002; Sinka et al., 2012). Elevated precipitation and humidity levels have also been shown to correlate with arboviral outbreaks by creating additional mosquito breeding sites and decreasing mosquito mortality, respectively (Harris et al., 2018; Hu et al., 2006; Li et al., 1985; Messina et al., 2016; Scott et al., 2000). However the exact relationships between these climate factors and vector suitability are often disputed and have yet to be rigorously established within in the field (Alto and Juliano, 2001; Canyon et al., 2013; Da Cruz Ferreira et al., 2017; Descloux et al., 2012; Naish et al., 2014).

Fitting epidemiological models to empirical data provides a way of quantifying these relationships. This has been done to successfully determine the local epidemiological drivers of Zika (Kucharski et al., 2016; Lourenço et al., 2017). Multiple facets of serological, spatial, phylogenetic or surveillance data can be linked to better elucidate the underlying mechanisms of disease, e.g. Kucharski et al. (2018); Wesolowski et al. (2015). Systems of ordinary differential equations have already been extensively fit to empirical data (Chowell et al., 2007; Lourenço et al., 2017; O’Reilly et al., 2018; Tuncer et al., 2018). However these deterministic frameworks fail to capture the inherent stochasticity and spatio-temporal heterogeneities of arboviral disease, and place strong implicit assumptions on vector ecology. Although discrete-time statistical transmission models (Li et al., 2018), such as Bayesian hierarchical dynamic Poisson models (Martínez-Bello et al., 2017), spatio-temporal risk models (Lowe et al., 2014; Martínez-Bello et al., 2018), and mixed models (Lowe et al., 2017), encapsulate the stochastic dynamics of arboviruses, they fail to capture the associations between epidemiological determinants and essential transmission drivers. Individual based models are arguably better suited to capture the spatio-temporal dynamics of arboviral disease whilst allowing for an unrestricted relationship between extrinsic and intrinsic factors.

Due to computational inefficiency, agent based frameworks have only previously been fit to surveillance data using a maximum likelihood approach, reporting only point estimates of each parameter of interest (Soda et al., 2018). Here, we demonstrate that the computational speed-up from GPU-acceleration allows us to fit a spatially-explicit, climate-driven individual based model to arboviral disease incidence data within a Bayesian framework. For this, we first fit our model to simulated data before considering empirical data from the first Zika outbreak in Brazil. This data was chosen because it has two clean disease outbreaks and as the primary vector for Zika and dengue are at least the same, dengue’s ecological drivers can be reasonably inferred. We also explore the effects of relaxing classical ordinary differential equation model assumptions of mosquito mortality rates and spatial dynamics on inference about mosquito ecology, and emphasise the importance of inter and intra-urban human mobility on vector-borne disease outbreaks. Finally, we show the forecasting potential of this framework, highlighting its potential usage as a real-time analysis tool for epidemiological outbreaks.

## 2 Methods and materials

### 2.1 Individual based model

Here, the individual based model as described by Lourenço and Recker (2013) was used, which included host and vector demography, an infection process and spatially-explicit community structure.

Within this model, age-dependent human, *µ*_*H*_, and mosquito mortality, *µ*_*V*_, was modelled from a bi-Weibull and Weibull distribution, respectively, defined by:

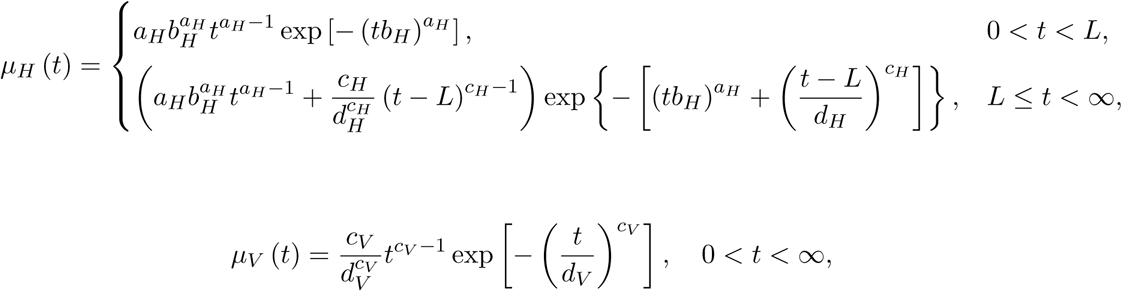

where *t* is the individual’s age in days, *L* is the location parameter of where infant and adult mortality coincide, *{a*_*H*_, *b*_*H*_*}* and *{c*_*H*_, *d*_*H*_*}* are the shape and scale parameters of infant and adult human mortality, respectively, and *{c*_*V*_, *d*_*V*_*}* are the shape and scale parameters of adult mosquito mortality, respectively.

An arbovirus was introduced into the system by allowing every human individual to be either susceptible (*S*_*H*_), exposed (*E*_*H*_), infectious (*I*_*H*_), or have recovered from (*R*_*H*_) the disease. Similarly, individual mosquitoes were either susceptible (*S*_*V*_), exposed (*E*_*V*_) or infectious (*I*_*V*_) with the virus. Mosquitoes bit at a constant rate *β* and infectious mosquitoes transmitted the virus with probability *p*_*H*_. Infected humans became infectious after 1*/E*_*H*_ days and recovered after 1*/γ* days. Mosquitoes were then infected with probability *p*_*V*_ given a bite on an infectious human, and became infectious after 1*/ϵ*_*V*_ days. To account for the introduction of the disease into the system, humans were also infected at an external infection rate, *ι*.

We modelled the influence of temperature, humidity and rainfall on these transmission parameters similarly to Lourenço et al. (2017).

#### 2.1.1 Temperature-dependent parameters

For the relationship between the extrinsic incubation period, 1*/ϵ* _*V*_, and temperature, *T,* (in Celsius) we applied the formulation by Focks et al. (1993) (see also Focks et al. (1995) and (Otero et al., 2006)), motivated by the enzyme kinetic model by Sharpe and DeMichele (1977), which assumed that replication is determined by a single rate-controlling enzyme. In order to match the daily time steps of the individual based model, the formula was multiplied by 24, as the replication rate given by Focks et al. (1995) is defined per hour:

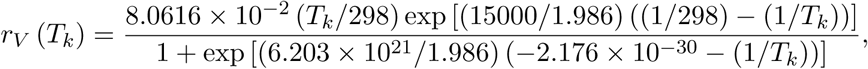

where *r*_*V*_ (*T*_*k*_) denotes the developmental rate of the virus in the mosquito at an environmental temperature *T*_*k*_ in degrees Kelvin. Under all environmentally realistic temperatures, the denominator can be approximated by one, thus we reduced the equation to:

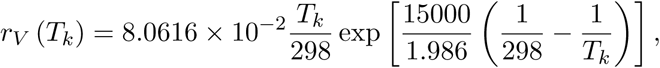

As the parameters in the temperature-dependent equation were estimated from laboratory experiments, where mosquitoes were infected with a fixed titre of virus, we scaled the replication rate linearly to account for the difference between infected blood meal virus titre in the laboratory versus in the field:

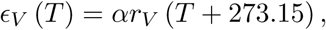

for some scalar *α >* 0 and *E*_*V*_ (*T)* represents the developmental rate of the virus in the mosquito at an environmental temperature *T* in degrees Celsius.

Arbovirus vector-to-human transmission probability, *p*_*V*_, generally increases with temperature but then sharply declines at very high temperatures (Lambrechts et al., 2011). This was described using a non-monotonic function given in the work by Mordecai et al. (2017) as:

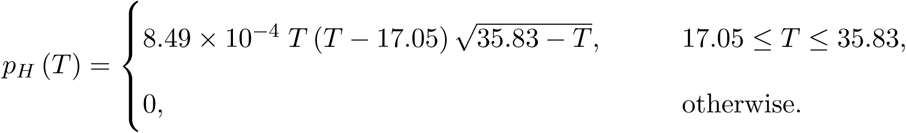

Similarly, the human-to-vector transmission probability was influenced by temperature (Mordecai et al., 2017) and given as:

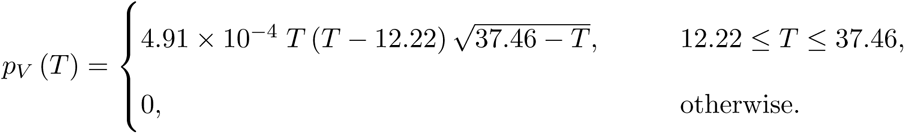

The relationship between temperature and mean life expectancy of a mosquito (1*/µ*_*V*_) was obtained using a fourth degree polynomial fit to data from a study by Yang et al. (2009):

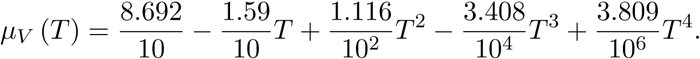

where *µ*_*V*_ (*T)* denotes the mortality rate of vectors at a temperature *T* in Celsius.

#### 2.1.2 Humidity-dependent parameters

Humidity, *Ĥ*, has a complex relationship with rainfall and temperature and is known to affect the birth and death rate of vectors. Humidity effects wee therefore modelled explicitly. Humidity was normalized between [0, 1] and then standardised about the mean, 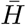, as follows (Tran et al., 2013):

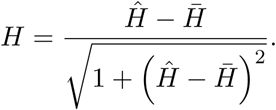

Therefore *H* ∈ [*-*1, 1]. The death rate of vectors was assumed to have a negative relationship with humidity (Alto and Juliano, 2001):

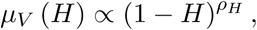

with some power *ρ*_*H*_ *>* 0 and *µ*_*V*_ (*H*) denotes the effect of humidity on the mosquito mortality rate. Combining this with the temperature-dependent effects on mosquito mortality gave,

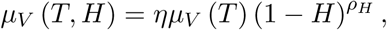

for some linear scalar *η >* 0. We then set *d*_*V*_, the scale parameter for the mosquito mortality distribution, equal to *µ*_*V*_ (*T, H*).

#### 2.1.3 Rainfall-dependent parameters

Increased rainfall ensures additional breeding sites resulting in increased rates of mosquito oviposition (Scott et al., 2000) and mosquito density. Rainfall, *R*, was smoothed using a moving average and then normalised, such that *R* ∈ [0, 1].

The expected change in the number of mosquitoes between time *t* and *t* + 1 (in days), denoted by Δ*N*_*V*_ (*t* + 1) was assumed to follow a logistic growth model with carrying capacity dictated by rainfall.

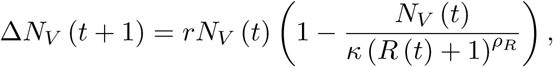

where *κ* is the minimum carrying capacity of the environment, *r* is the maximum growth rate and *ρ*_*R*_ *>* 0 scales the carrying capacity with rainfall.

#### 2.1.4 Spatial structure

We incorporated spatial structure by dividing the host and vector populations into a set of communities, *C*, which were then organised into a non-wrapping lattice. Humans and mosquitoes were homogeneously distributed throughout the lattice, and individuals were assumed to mix homogeneously within each community. We then allowed infection events from each community to disperse:

i. locally, where infectious individuals can infect those in surrounding communities with probability 1 *-p*_*σ*_, and
ii. across long distances, where infectious individuals can infect anyone within the lattice with probability *ω*.

#### 2.1.5 Computational implementation

In order to alleviate the extreme computational costs of an individual based model we implemented the model within a GPU-accelerated framework. This achieved massive speed-ups (Table 1) which then allowed us to consider fitting the individual based model within a fully Bayesian framework for the first time.

**Table 1.** Simulation run-times on the CPU versus GPU. The simulation using both the CPU and GPU-accelerated implementations of the model for 100 simulation years, with 3.5 million humans and 7 million mosquitoes. The CPU implementation of the model was parallelised on 8 threads and executed on an Intel Core i7-4790K CPU @ 4.00Ghz processor. The GPU implementation of the model was both executed using an NVIDIA GeForce GTX 970 2GB graphics card and an NVIDIA GeForce GTX 1080 Ti 11GB graphics card.

| Parallelisation | Runtime | Speed-up |
| --- | --- | --- |
| CPU | 40 minutes | - |
| GPU with GTX 970 | 1 minute | 40 |
| GPU with GTX 1080Ti | 25 seconds | 96 |

#### 2.1.6 Model parameters

Unless stated otherwise, the parameters used during individual based model fitting are shown in Table 2. We fixed these parameters values to best represent the epidemiology of arboviral disease within an urban setting as informed by the literature. The parameters that were inferred during model fitting are shown in Table 3. We sought to estimate these unobserved parameters as they are not currently well established.

**Table 2.**
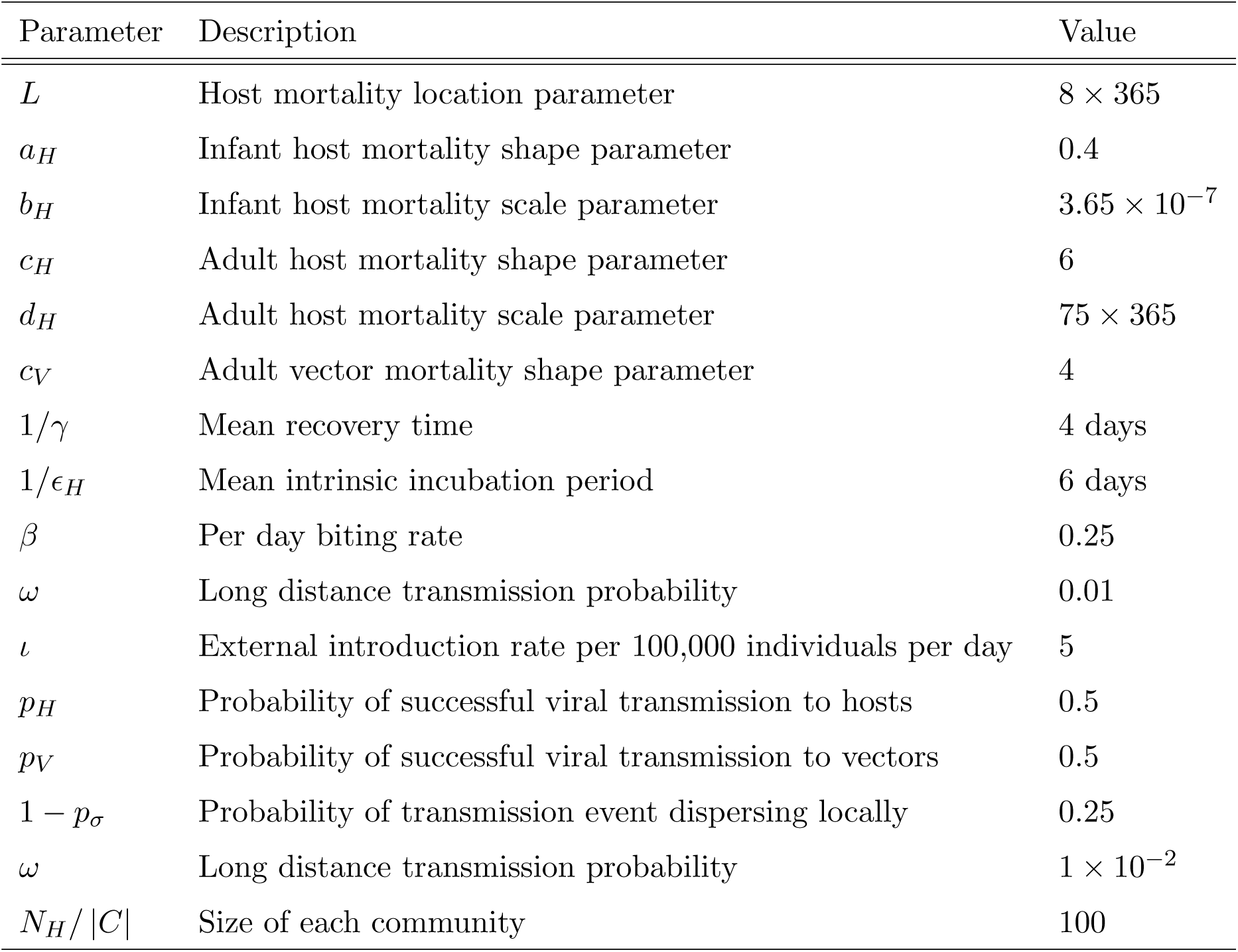
The default set of fixed parameter values used in fitting the individual based model to weekly Zika incidence from 2015–2017 in Feira de Santana, Brazil.

**Table 3.** Unobserved parameters to be estimated by the Metropolis-Hastings MCMC algorithm in conjunction with the individual based model and given climate and incidence data.

| Unobserved parameter | Description | Range |
| --- | --- | --- |
| $\alpha$ | Scalar for extrinsic incubation period | $(0, \infty)$ |
| $\eta$ | Scalar for mean adult vector life expectancy | $(0, \infty)$ |
| $\rho_R$ | Non-linear scalar for rainfall seasonality | $(0, \infty)$ |
| $\rho_H$ | Non-linear scalar for humidity seasonality | $(0, \infty)$ |
| $\kappa$ | Minimum mosquito carrying capacity | $(0, \infty)$ |
| $\phi$ | Reporting dispersion | $(0, 1)$ |
| $p_{obs}$ | Observation rate | $(0, 1)$ |

### 2.2 Bayesian specification

We specified the model in a Bayesian framework to determine the posterior distribution, *π* (*θ*|*y*), the set of unobserved parameters, *θ*, (given in Table 3) given outbreak data *y*, such that,

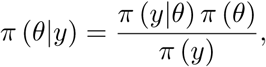

where *π* (.) represents a probability distribution, *π* (*y*|*θ*) denotes the likelihood distribution of the data *y* given the parameters *θ, π* (*θ*) is the prior distribution of *θ*, and *π* (*y*) is the marginal likelihood of the data *y*, specified by,

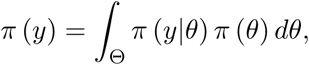

where Θ denotes the multi-dimensional space of all possible values of *θ*.

### 2.3 Markov chain Monte Carlo model fitting

For a complex model, the posterior distribution of parameters *θ* cannot be written in a closed form as the integral *π* (*y*) is impractical to evaluate analytically. Instead, we fit the model using Markov chain Monte Carlo (MCMC) methods. A Markov chain is a sequence of stochastic events where the probability of each event occurring depends only upon the state of the previous event. Under certain conditions, a Markov chain can be set up such that it converges to a stationary distribution, and thus once converged, the chain produces random samples from the stationary distribution. This allowed us to take a large number of samples from the posterior distribution for the parameters through construction of a Markov chain that converges to the desired posterior distribution. From this, we could estimate the posterior mean and credible intervals of the parameters of interest. The classical Metropolis-Hastings MCMC algorithm that was employed here is defined in Algorithm 1 (Hastings, 1970; Metropolis et al., 1953).

#### Algorithm 1: Random-walk Metropolis-Hastings MCMC

Let *θ*_*k*_ denote the set of parameters at position *k ∈* ℕ_0_ in the Markov chain, then the next set of unobserved parameters in the chain, denoted *θ*_*k*+1_, is generated as follows:

1. Sample a candidate parameter set *θ′* from some proposal distribution with probability density function *q* (.*|θ*_*k*_).
ii. Compute the acceptance ratio of the proposed parameter set *θ*′ given the previous parameter set *θ*_*k*_:

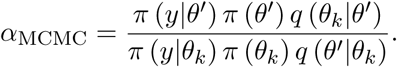
iii. Generate *u ∼* Unif (0, 1).
iv. If *u < α*_MCMC_, accept the candidate parameters and set *θ*_*k*+1_ = *θ*′, else reject the candidate parameter set and use the previously accepted parameter set, thereby setting *θ*_*k*+1_ = *θ*_*k*_.

Here *π* (*y|θ*) is the likelihood of the case data *y* given a parameter set *θ, π* (*θ*) is the joint-prior distribution of all unobserved parameters, and *q* (.) is the probability density function of the proposal distribution of *θ*.

#### 2.3.1 Likelihood

Let *y* denote the time series of the number of weekly infected cases in the observed data. The likelihood of the data *y* given parameters *θ* is often dependent on some unobserved, or hidden, variables, which we denote *z* and corresponds to the time series of the actual number of weekly infected cases, where *y* is the number of those actual cases which are observed. Therefore, the likelihood can be written as,

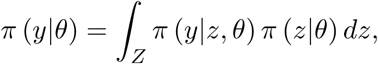

where *Z* is the multidimensional space containing all possible values of *z*. Here, *π* (*y*|*z, θ*) represents an observation process from the time series of the actual number of weekly infected cases, *z*, to the observed data *y*, and *π* (*z*|*θ*) denotes the probability density of the unobserved states *z* given parameters *θ*. As this integral is intractable in practice, we approximate the integral using importance sampling, such that for a given number of simulated data sets, *N,* we can write the following unbiased estimator for the integral (Andrieu and Roberts, 2009; Beaumont, 2003):

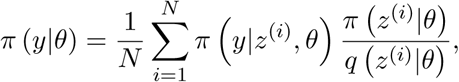

where *z*^(*i*)^ denotes the *i*-th replicate from the model and *q z*^(*i*)^|*θ* denotes the probability density of the importance sampling distribution. By simulating from the underlying individual based model, we have *π* (*z*^(*i*)^|*θ*)= (*q z*^(*i*)^|*θ*), and so the estimated likelihood became:

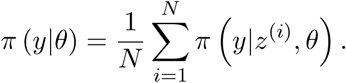

Each data set was a weekly time series, and the observation process was assumed independent given the hidden states *z*, therefore,

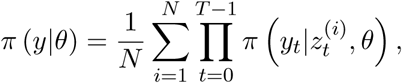

where *T* is the maximum number of weeks in the data, *y*_*t*_ is the incidence at week *t* in the observed data, and 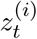 is the incidence in week *t* of the *i*-th replicate simulated from the model. Finally, we assumed that the observation process is negatively binomially distributed with some observation rate *p*_*obs*_ and dispersion parameter *ϕ* to account for under-reporting of the true incidence:

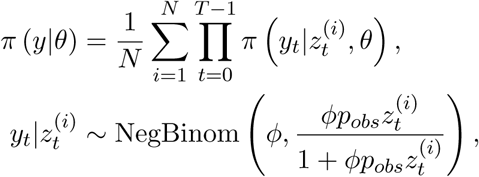

where the negative binomial distribution was parametrised with mean 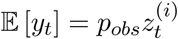 and variance 𝔼[*y*_*t*_] + *ϕ* 𝔼[*y*_*t*_]^2^.

It has been shown that substitution of an unbiased estimate of *π* (*y*|*θ*), such as above, into the classical Metropolis-Hastings MCMC algorithm, Algorithm 1, produces samples from the true posterior distribution of *y* given *θ* in probability (Andrieu and Roberts, 2009).

#### 2.3.2 Prior distributions

Weakly informative priors were inferred from field and laboratory experiments (for Zika and dengue), the parametrization of our simulation model, and previous model fits from the scientific literature. These are listed in Table 4.

**Table 4.** The prior of each parameter used during fitting the stochastic individual based model to outbreak incidence data, where TruncNorm (*µ, Σ, a, b*) denotes a truncated normal distribution with mean *µ*, standard deviation *Σ*, lower bound *a*, and upper bound *b*.

| Parameter | Prior | References |
| --- | --- | --- |
| $\alpha$ | $\text{TruncNorm}(2.5, 0.5, 0, \infty)$ | Chan and Johansson (2012); Krow-Lucal et al. (2017) |
| $\eta$ | $\text{TruncNorm}(3, 0.5, 0, \infty)$ | Maciel-de Freitas et al. (2007); Marinho et al. (2016); Muir and Kay (1998) |
| $\rho_R$ | $\text{TruncNorm}(1, 0.4, 0, \infty)$ | - |
| $\rho_H$ | $\text{TruncNorm}(1, 0.4, 0, \infty)$ | - |
| $\kappa$ | $\text{TruncNorm}(3, 1, 0, \infty)$ | - |
| $\phi$ | $\text{Beta}(20, 80)$ | Kucharski et al. (2018) |
| $p_{obs}$ | $\text{Beta}(10, 90)$ | Kucharski et al. (2016); Lourenço et al. (2017); Shutt et al. (2017) |

#### 2.3.3 Proposal distribution

Candidate parameters *θ*′ were sampled from a multivariate normal distribution with probability density function *q* (.) using a random-walk adaptive routine as detailed in Roberts and Rosenthal (2009). The routine uses the existing covariance structure in the Markov chain of *θ* to adapt toward an optimal proposal distribution that promotes well-mixing chains. Using the optimal scaling parameter defined in Sherlock et al. (2015), we define the proposal distribution here as,

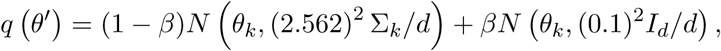

where *θ*_*k*_ is the set of parameters at the *k*-th position in the chain, Σ_*k*_ is the current estimate of the covariance structure of the posterior distribution, defined as the empirical covariance matrix from the current samples for *θ* at iteration *k, d* is the dimension of Σ_*k*_, *β* is a small positive constant (here, we take *β* = 0.05), and *I*_*d*_ is the identity matrix of dimension *d*.

#### 2.3.4 Convergence

In order to allow the chains to converge to the posterior distribution, unless stated otherwise, the first 20,000 iterations of each model fit were discarded as burn-in. Furthermore, in order to assess the independence of parameter initialisation on the convergence of the model, each unobserved parameter in the Markov chain was initialised from multiple randomly sampled points of each prior. Convergence to the same posterior distribution was then assessed through visual inspection of the Markov chains. Thereafter, each parameter in the Markov chain was initialised from the mean of each prior.

#### 2.3.5 Optimisation

The likelihood of the data *y* given parameters *θ* depended upon the number of simulated data sets, *N,* generated per proposed parameter set. At the cost of computational runtime, a higher number of simulations per step increases decreased the variance in the estimated likelihood, *π* (*y*|*θ*), and so the acceptance rate of the algorithm increased and in turn, the mixing chains improved. To balance computational cost with how well-mixed each of the parameters were, *N* was selected such that the variance of the log of the estimate of the marginal likelihood was approximately equal to one (Doucet et al., 2015; Pitt et al., 2012; Sherlock et al., 2015). As a result, unless stated otherwise, the number of simulated time-series per step of the MCMC, *N,* was set to be equal to thirty.

### 2.4 Incidence and climate data

The presented climate and weekly notified case data of Zika in Feira de Santana, Brazil from 1st February 2015 to 31st December 2016 were taken from Lourenço et al. (2017).

## 3 Results

As a proof of concept, we first fit the model to simulated data from the model itself, before fitting to weekly notified case data of Zika in Feira de Santana. Because there was only one circulating serotype during the first Zika outbreak in Brazil, both cases only considered a single serotype and a fully susceptible host population.

### 3.1 Model fit to simulated incidence data

In order to demonstrate that the unobserved parameters can be inferred from relatively sparse data, the model was first fit within a Bayesian framework to incidence data simulated from the individual based model with the preselected unobserved parameter values listed in Table 5. The incidence data exhibited two disease outbreaks, one in the middle of the first year, peaking with high rainfall and humidity levels, and the other at the start of the second year (Figure 1).

**Table 5.** Values of the unobserved parameters used to generate (simulated) weekly case data.

| Unobserved parameter | Description | Value |
| --- | --- | --- |
| $\alpha$ | Scalar for extrinsic incubation period | 3 |
| $\eta$ | Scalar for mean adult vector life expectancy | 2 |
| $\rho_R$ | Non-linear scalar for rainfall seasonality | 1.5 |
| $\rho_H$ | Non-linear scalar for humidity seasonality | 0.5 |
| $\kappa$ | Minimum mosquito carrying capacity | 2.3 |
| $\phi$ | Reporting dispersion | 0.05 |
| $p_{obs}$ | Observation rate | 0.2 |

**Figure 1.**
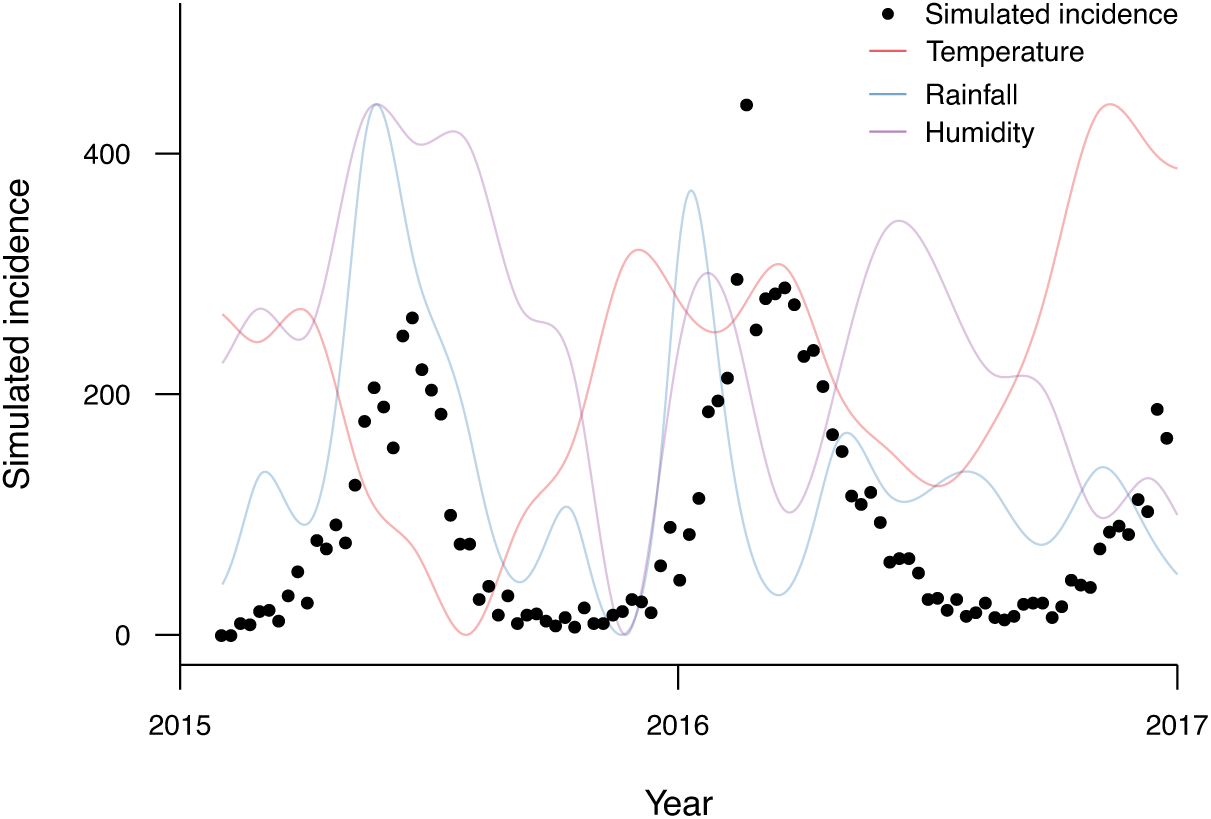
Observed (simulated) incidence and climate dynamics. Together with climate data from Feira de Santana, the individual based model was executed on preselected unobserved parameter values shown in Table 5, resulting in two disease outbreaks over two years.

Model generated observations matched the temporal signature of the original incidence data (Figure 2A). There was large uncertainty around the simulated dynamics, however this was not due to the stochasticity of the individual based model itself, but due to the underlying observation process. That is, we selected an overly-dispersed observation process, which created high variance between credible outcomes in observed incidence during both epidemic peaks. In contrast, the variance in the estimated percentage of the population that were affected during the first year of the outbreak, or 2015 attack rate, was small, with the true attack rate of the original simulated data contained within the 95% credible interval of the attack rate (Figure 2B).

**Figure 2.**
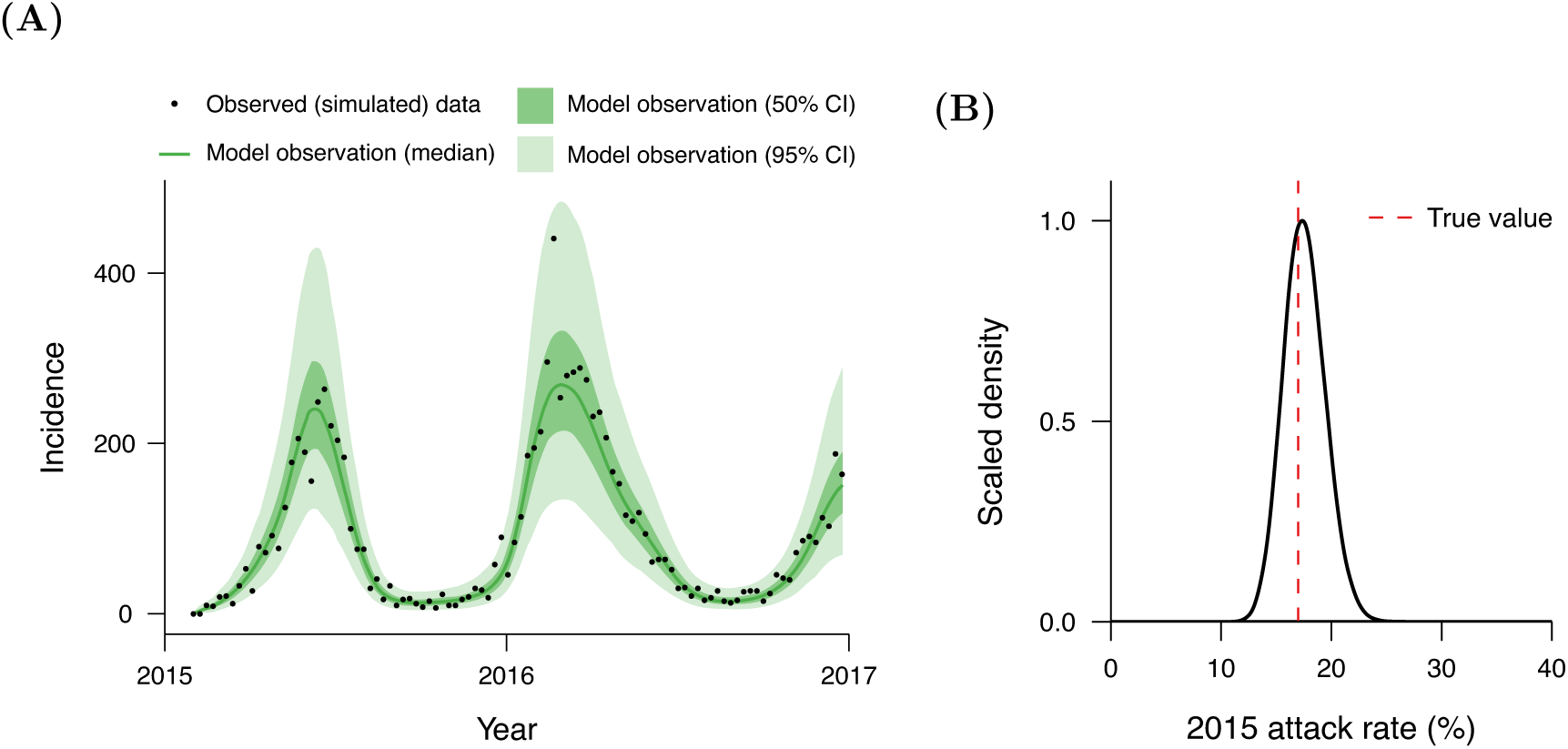
Posterior predictive distribution and attack rate of simulated data. **(A)** The temporal dynamics of the distribution of simulated observed cases exhibited similar underlying dynamics to the incidence data generated from the individual based model. **(B)** The inferred percentage of the total population infected during the first year contained the true attack rate of the simulated data. The posterior predictive distribution was calculated from randomly sampling 1,000 sets of parameter values from the posterior distribution and simulating these within the individual based model. Weekly observed cases are then randomly sampled from the negative binomial distribution, representing the observation process from total to notified cases, using the dispersion parameter, *ϕ*, mean probability of observing a single case, *p*_*obs*_, and simulated total weekly cases from the individual based model.

These findings demonstrated that the individual based model could accurately fit to incidence data. However, fitting to this data came as no surprise, as a model with so many parameters is likely to give a good fit. What was important here, were the inferred unobserved parameter values themselves.

#### 3.1.1 Posterior distribution

The 95% credible intervals of the posterior distribution of each unobserved parameter also included the true values used to generate the incidence data to which the individual based model was fit (Figure 3). We found that the true parameter values for the scalars for the extrinsic incubation period, mosquito life expectancy and observation rate were well-inferred, however true values for the minimum mosquito-to-human ratio and rainfall and humidity effects were relatively unlikely. The inferred over-dispersion parameter was greater than the true value because estimating the overdispersion parameter of only approximately 100 samples is generally challenging. The high uncertainty in the minimum mosquito-to-human ratio and scalars for rainfall and humidity effects was likely a result of strong correlations between the inferred values for each.

**Figure 3.**
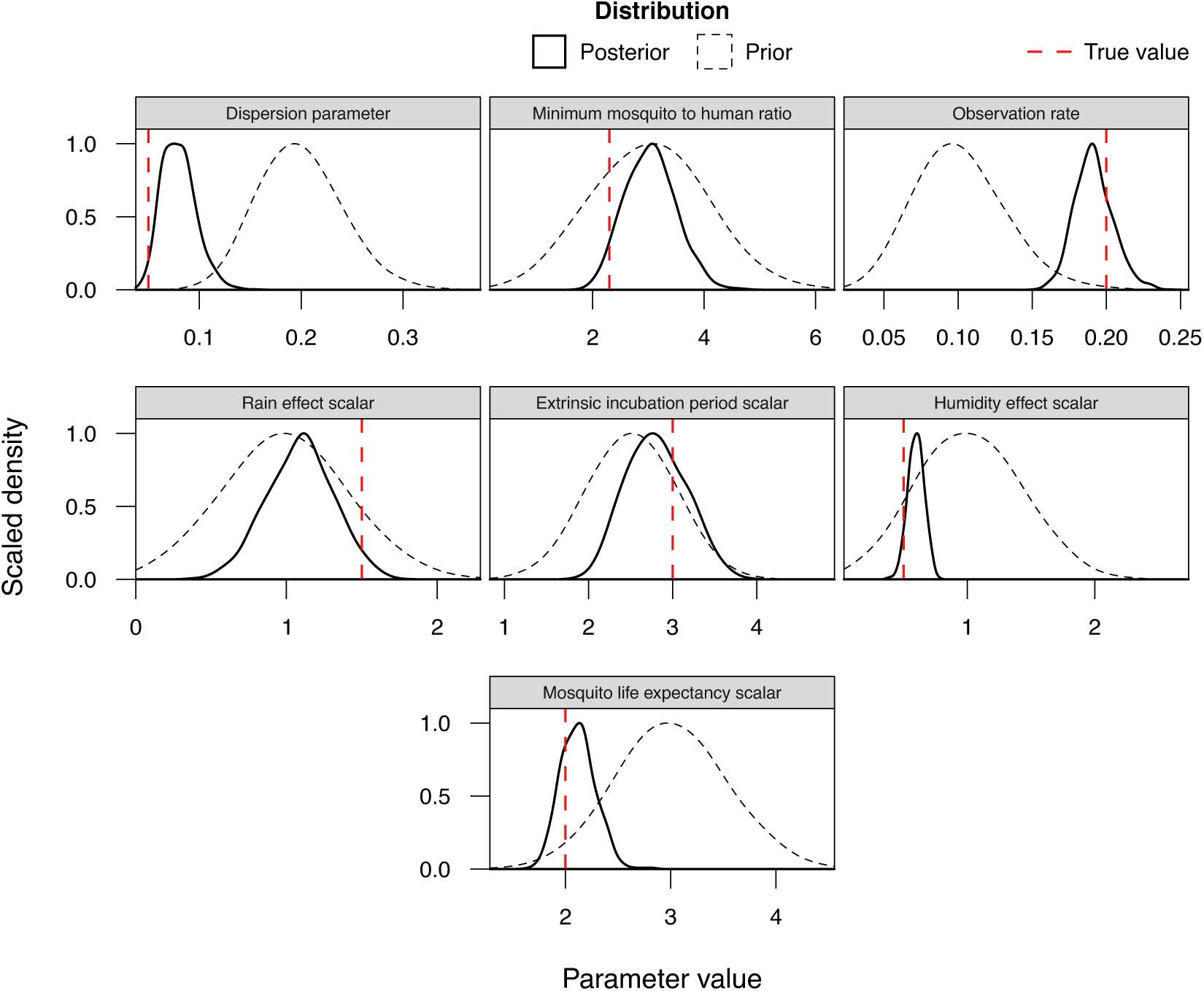
Posterior distributions of unobserved parameters. All estimated posterior distributions from the fitting process were unimodal and improved upon the weakly informative prior distributions. The inferred unobserved parameters were the dispersion parameter for the observation process that mapped total weekly cases to notified cases, *ϕ*, the minimum mosquito-to-human ratio, *κ*, the probability of a case being notified, *p*_*obs*_, the effect of rainfall on mosquito density, *ρ*_*R*_, the scalar that influences the extrinsic incubation period, *α*, the effect of humidity on mosquito longevity, *ρ*_*H*_, and the scalar that controls mosquito mortality rates, *η*. Here, posterior distributions were calculated from chains of 125,000 iterations with a burn in period 20,000 iterations.

#### 3.1.2 Correlations of unobserved parameters

There was a strong negative correlation (Pearson’s *r* = *-*0.68) between the minimum mosquito-to-human ratio and the effect of rainfall on the mosquito-to-human ratio (Figure 4A). In order to achieve the same transmission potential during outbreaks, smaller baseline values of mosquito density required an increasing effect of rainfall so that the mosquito density during outbreaks is maintained. Similarly, there was a strong positive correlation (*r* = 0.62) between the minimum mosquito-to-human ratio and the effect of humidity on mosquito longevity (Figure 4B). This was because lower mosquito densities throughout periods of low transmission required increased mosquito longevity in order sustain sufficient transmission potential.

**Figure 4.**
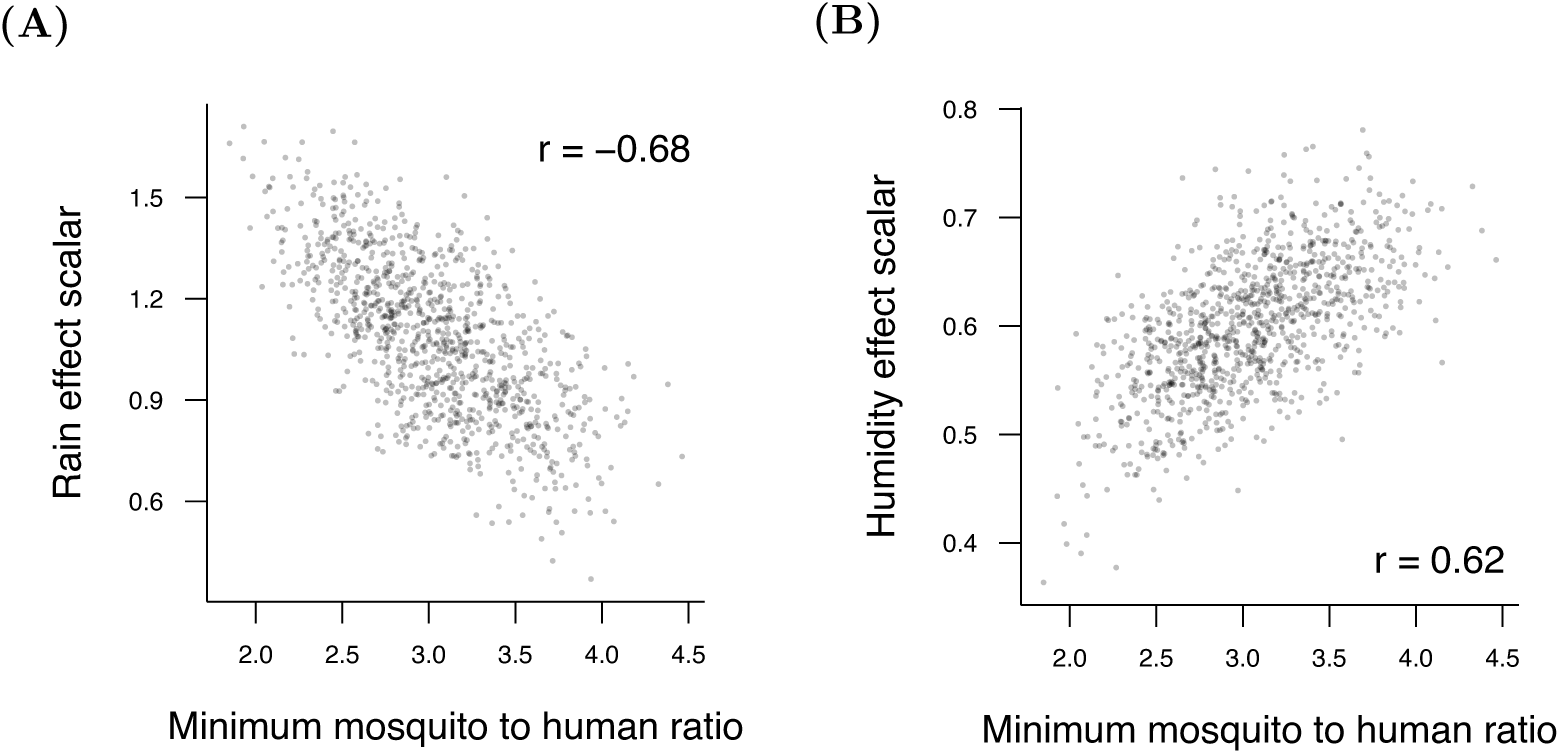
Correlation between rainfall, humidity and the minimum mosquito-to-human ratio. **(A)** There was a strong negative correlation between the effect of rainfall on mosquito density and the minimum mosquito-to-human ratio. **(B)** Similarly, there was a strong positive correlation between the minimum mosquito-to-human ratio and the effect of humidity on mosquito longevity. Pearson’s correlation coefficients, *r*, shown were calculated from 2000 samples from the posterior distribution.

#### 3.1.3 Key transmission parameters were well-inferred

To demonstrate the relationships between key epidemiological parameters, the posterior distributions of these parameters were translated into distributions of the mosquito-to-human ratio and mosquito life expectancy at the peak of the first epidemic. The inferred estimates of the human to mosquito ratio and mosquito life expectancy during the first outbreak were well-inferred, with the true values of each contained within the 95% credible intervals of each distribution (Figure 5).

**Figure 5.**
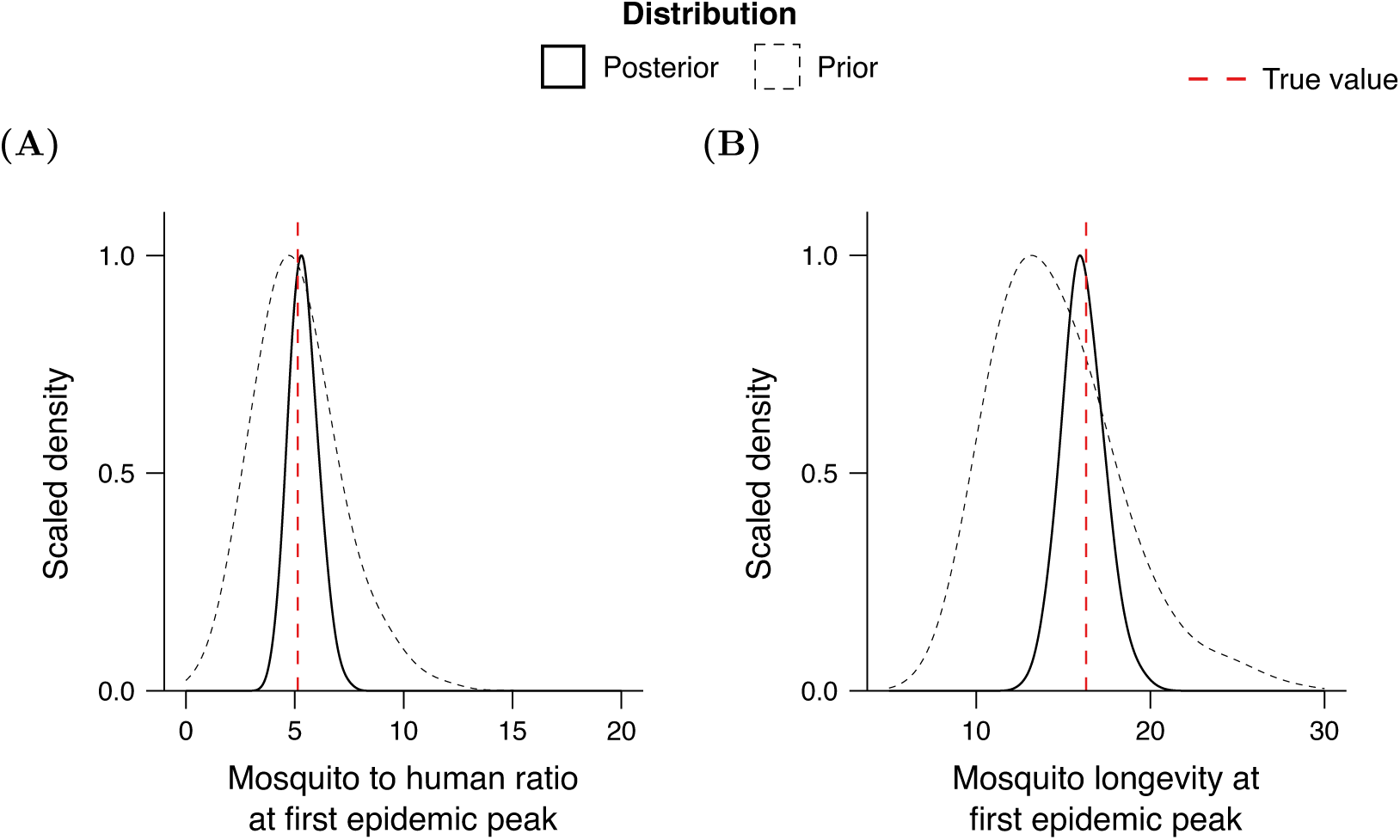
Key transmission parameters can be well-inferred from sparse data. By fitting the modelling framework to incidence data that was generated from the individual based model itself, the posterior distributions for **(A)** the mosquito-to-human ratio and **(B)** mean mosquito life expectancy contained the true parameter values used to generate the incidence data.

### 3.2 Model fit to empirical incidence data

From February 2015 to December 2016, Feira de Santana experienced two outbreaks of Zika: a large epidemic from April 2015 to August 2015 when rainfall and humidity levels were at their maximum over the two year period, and a small outbreak at the start of 2016 when temperature, humidity, and rainfall were consistently high (Figure 6). Now that we have shown that the model can capture the temporal dynamics of simulated incidence data, the individual based model was fit to weekly notified cases of Zika in Feira de Santana, Brazil.

**Figure 6.**
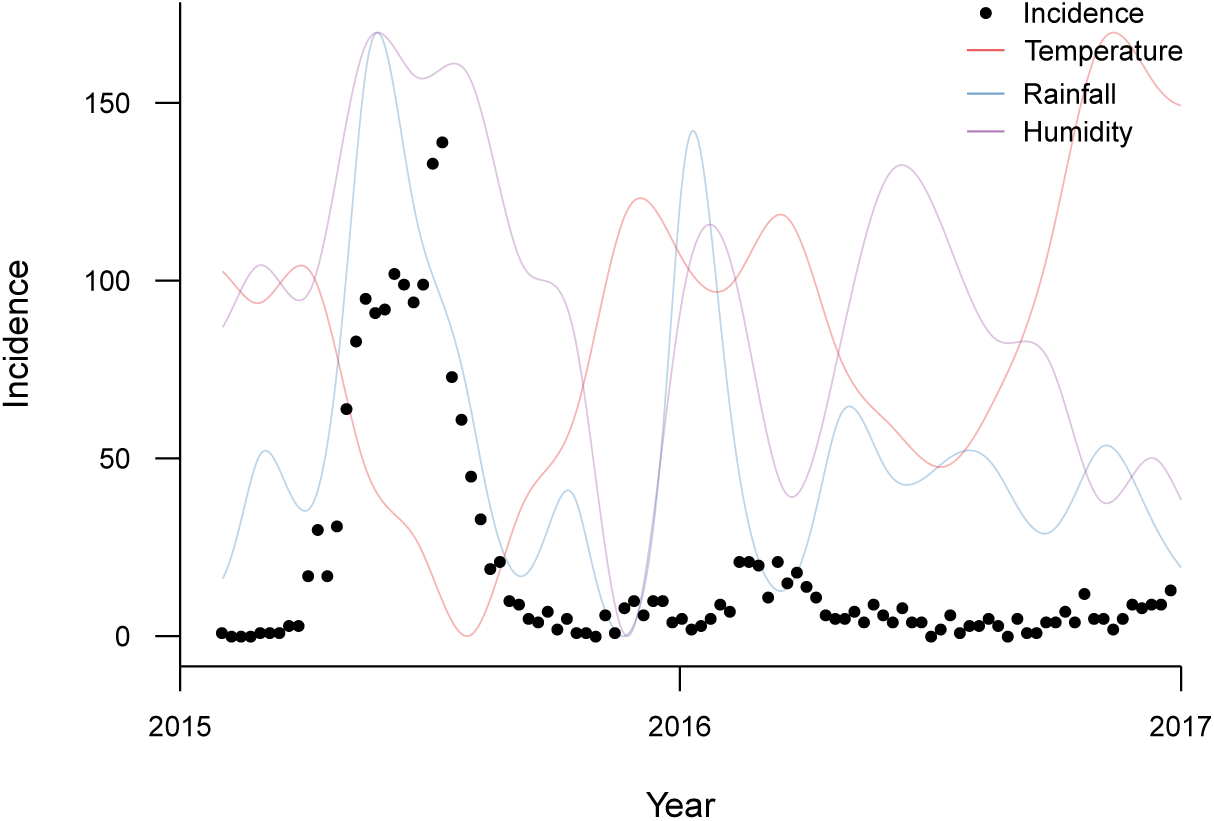
Observed incidence and climate dynamics. The individual based model was fit to weekly notified cases of Zika in Feira de Santana, Brazil between February 2015 and December 2016. There was one large epidemic during 2015 and a smaller outbreak at the start of 2016, both which correlate with elevated levels of rainfall and humidity, and the latter with high temperatures.

Fitting the model yields simulated dynamics that behave similarly to the dynamics of the empirical data, with a large epidemic from April 2015 to August 2015, and a much smaller outbreak at the start of 2016 (Figure 7A). There were high levels of uncertainty in the distribution of model generated observed cases during the 2015 outbreak because of the large variance in the underlying observation process. Furthermore, the percentage of the total human population infected, or attack rate, of around 50% during 2015 (Figure 7B) with very large uncertainty, between 20% and 70%.

**Figure 7.**
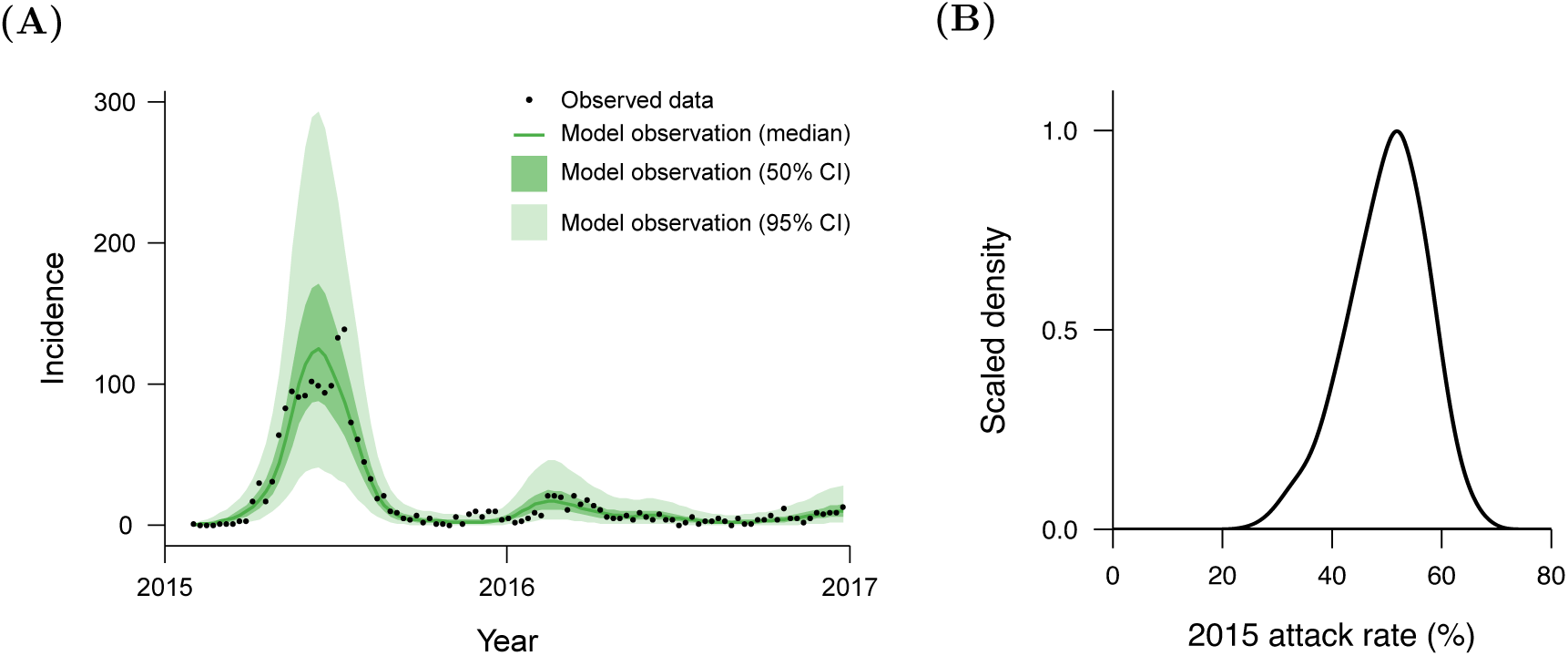
Posterior predictive distribution and attack rate. **(A)** The temporal dynamics of the distribution of simulated observed cases exhibited similar underlying dynamics to the weekly notified cases for Zika in Feira de Santana, Brazil, between 2015 and 2017. **(B)** The percentage of the total population infected during 2015 was inferred to be around 50%. The posterior predictive distribution is calculated from randomly sampling 1,000 sets of parameter values from the posterior distribution and simulating these within the individual based model. Weekly observed cases are then randomly sampled from the negative binomial distribution, representing the observation process from total to notified cases, using the dispersion parameter, *ϕ*, mean probability of observing a single case, *p*_*obs*_, and simulated total weekly cases from the individual based model.

The chains of accepted unobserved parameter values were well-mixed (Figure 8) and converged in probability to the posterior distributions of each unobserved parameter. All posterior distributions were unimodal, and the majority of posteriors narrowed from the weakly informative prior distributions that were set (Figure 9). The dispersion parameter, *ϕ*, for the negative binomial observation process from total weekly cases to notified weekly cases was slightly lower than expected, but exhibited a high degree of uncertainty. The posterior for the minimum mosquito-to-human ratio, *κ*, constricted to values between 2.0 and 6.0, as a sufficiently high baseline level of mosquito density was required for out of season transmission. Interestingly, the posterior distribution of the observation rate, *p*_*obs*_, greatly shifted and narrowed from the prior to values of less than 5%. The posterior distributions for the scalar that influences mosquito mortality rates, *η*, also exhibited a strong shift towards lower values from the prior distribution, and thus in the direction of longer mean mosquito life expectancy. In contrast, the scalar that controls the replication rate of the virus, *α*, moved towards higher values from the prior distribution, resulting in shorter extrinsic incubation periods.

**Figure 8.**
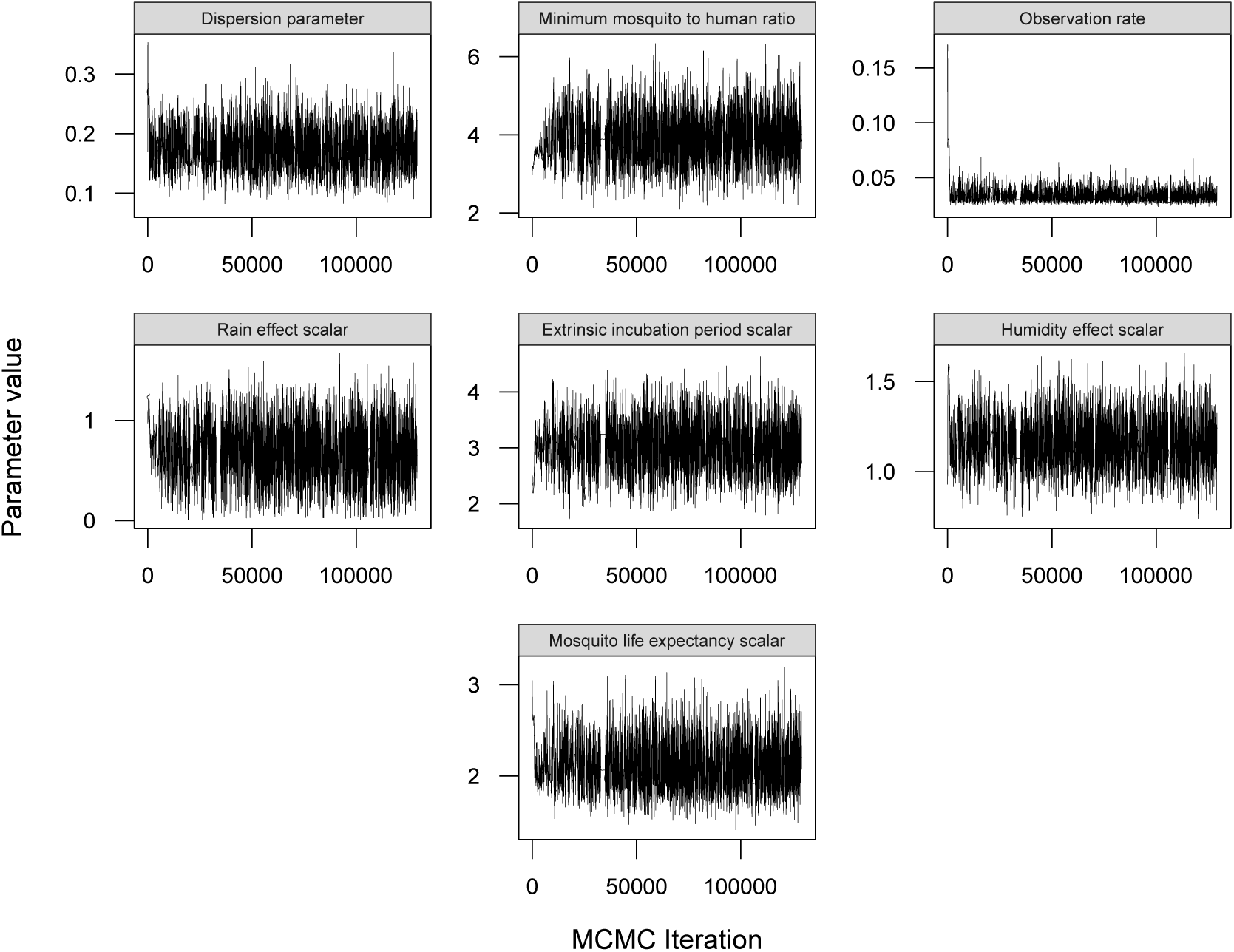
Markov chain Monte Carlo (MCMC) chains. Fitting the individual based model with the pseudo-marginal method with *N* = 30 particles per iteration to the empirical data yielded chains of accepted unobserved parameter values that were well-mixed.

**Figure 9.**
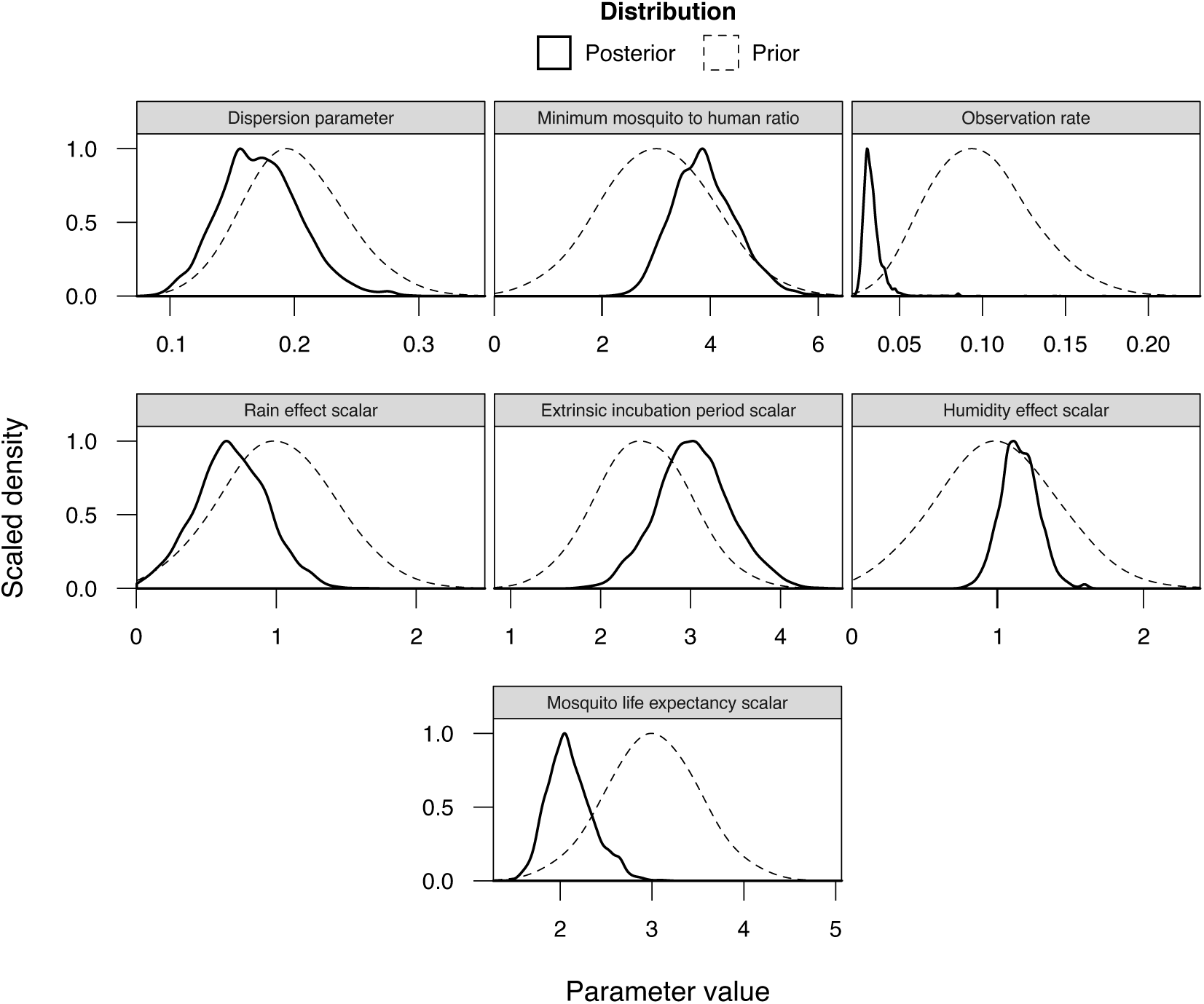
Posterior distributions of unobserved parameters. All estimated posterior distributions from the fitting process were unimodal and improved upon the weakly informative prior distributions. The inferred unobserved parameters were the dispersion parameter for the observation process that mapped total weekly cases to notified cases, *ϕ*, the minimum mosquito-to-human ratio, *κ*, the probability of a case being notified, *p*_*obs*_, the effect of rainfall on mosquito density, *ρ*_*R*_, the scalar that influences the extrinsic incubation period, *α*, the effect of humidity on mosquito longevity, *ρ*_*H*_, and the scalar that controls mosquito mortality rates, *η*. Here, posterior distributions were calculated from chains of 125,000 iterations with a burn in period 20,000 iterations.

Fitting the model to the empirical data suggests that the effect of rainfall on mosquito population density, *ρ*_*R*_, was weak (less than a linear response). This means that rainfall only caused small amplitude oscillations in mosquito density. However due to the association of high rainfall levels with large numbers of reported cases, the maximum number of mosquitoes per human was also found to be highest at peak incidence (Figure 10A). In contrast, the strong effect of humidity on mosquito longevity yielded pronounced oscillations in mean mosquito life expectancy that also correlated with incidence (Figure 10B). However, the relatively low temperatures during the 2015 outbreak modulated mosquito longevity from extremely high values. The extrinsic incubation period was also found to be consistently low throughout the study period, maximising during the 2015 epidemic due to relatively low temperatures (Figure 10C).

**Figure 10.**
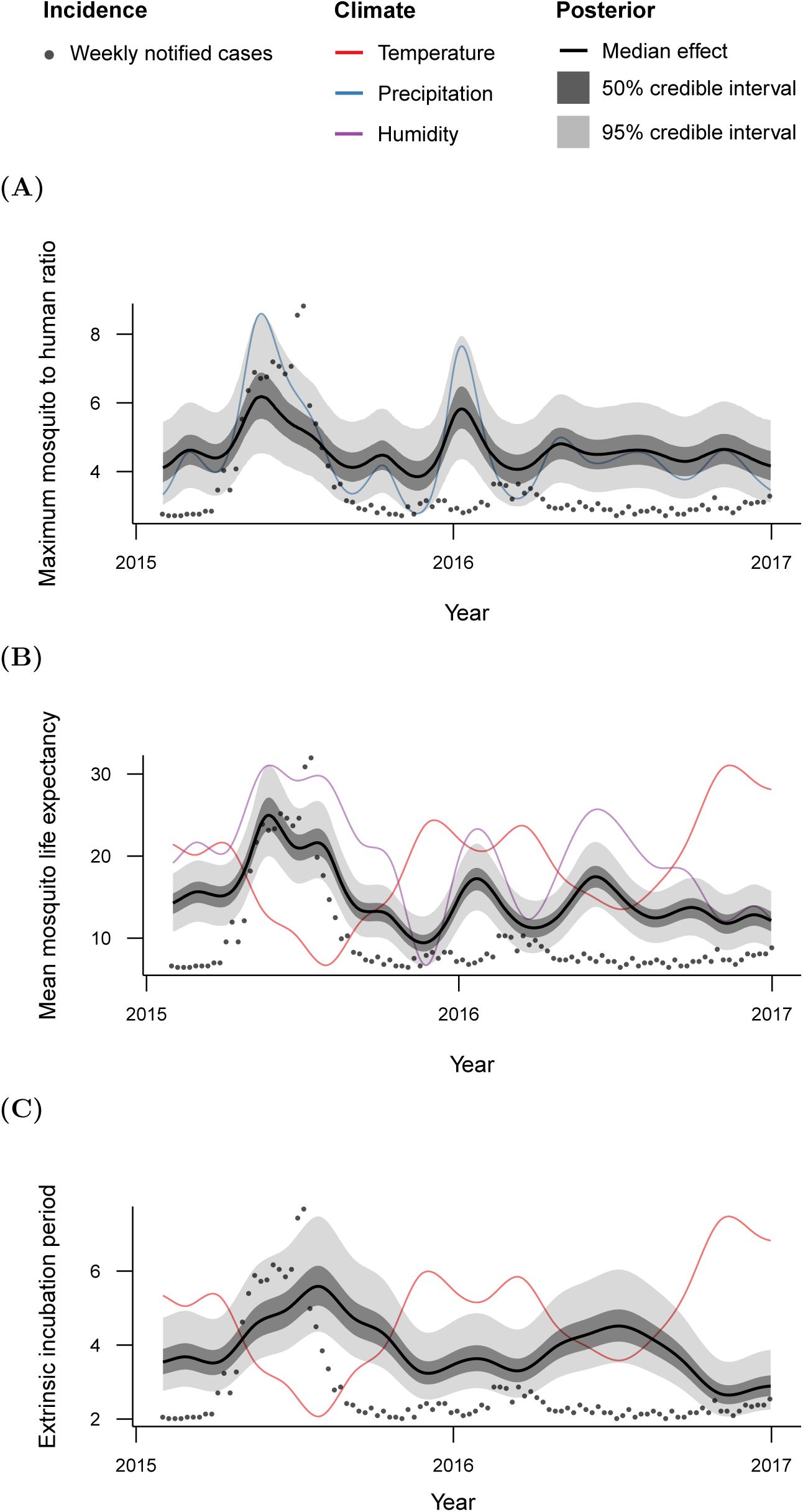
Inferred temporal dynamics of transmission parameters. Combining the inferred posterior distributions of unobserved parameters with the relationships defined between climate parameters and key transmission parameters, yield posterior distributions of each transmission parameter over time. During the 2015 outbreak, there was high **(A)** mosquito longevity, **(B)** mosquito density and **(C)** extrinsic incubation periods. During the periods of low transmission, extrinsic incubation periods, mosquito longevity and mosquito density were inferred to be consistently low.

Similarly to fitting to the simulated data, we also found correlations between some of the inferred unobserved parameters (Figure 11). The minimum mosquito-to-human ratio, *κ*, was found to be moderately negatively correlated with the effect of rainfall on mosquito population density, *ρ*_*R*_, with a Pearson’s correlation coefficient of *r* = −0.54. Lower mosquito densities required an increased influence of rainfall on mosquito density in order to maintain sufficiently high maximum mosquito-to-human ratios, and thus transmission potential, during periods of high transmission. Furthermore, the minimum mosquito-to-human ratio was positively correlated (*r* = 0.5) with the mosquito life expectancy scalar, *η*. This was because in order to sustain transmission potential, lower mosquito densities need to be counterbalanced by higher mosquito life expectancies, gained through lower scalars for mosquito longevity. The effect of rainfall on mosquito density was negatively correlated (*r* = −0.5) with the effect of humidity on mosquito longevity, *ρ*_*H*_, due to the strong positive correlation between the temporal dynamics of rainfall and humidity (*r* = 0.73). In line with the previous results, there was a moderate positive correlation (*r* = 0.46) between the effects of humidity on mosquito longevity and the scalar for mosquito mortality rates. Amplified oscillations in mosquito longevity need to be offset by a reduction in mean mosquito longevity, and thus an increase in the corresponding scalar, to maintain high transmission during the 2015 outbreak. Expectedly, the scalar influencing mosquito mortality rates was positively correlated with the scalar controlling the extrinsic incubation period (*r* = 0.34) as shorter mosquito life expectancy compensates for shorter extrinsic incubation periods, keeping the mean duration of infectivity in mosquitoes consistent. Finally, the mean probability of observing an infected human case, *p*_*obs*_, was strongly positively correlated with the effect of humidity on mosquito longevity (*r* = 0.58) and the scalar controlling mosquito longevity (*r* = 0.64). Increased humidity effects and scalers for mosquito longevity decreased the mean mosquito life expectancy throughout the study period. Therefore, the overall reduction in transmission potential was compensated for by increased observation rates.

**Figure 11.**
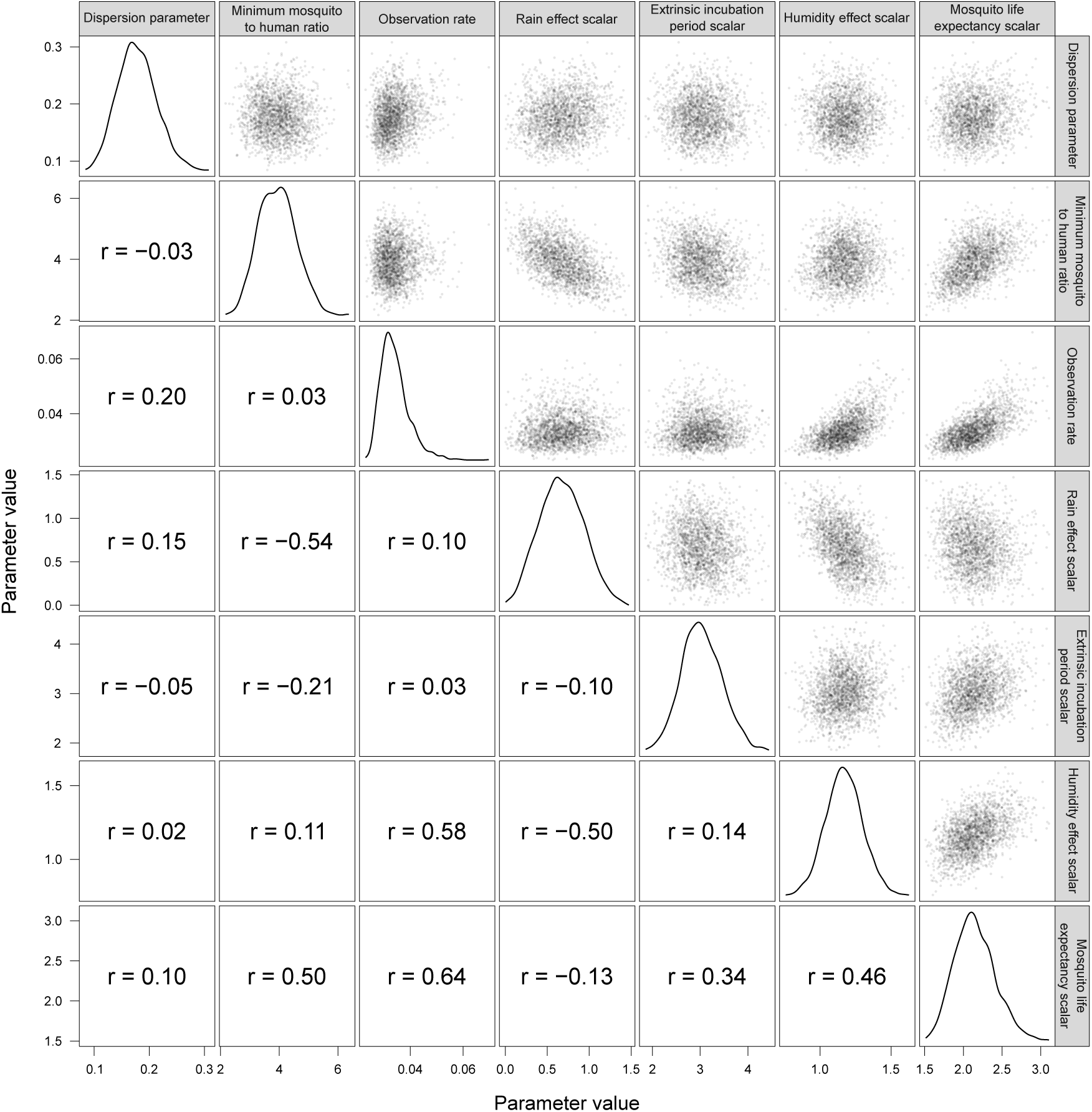
Correlations between accepted unobserved parameters. Moderate correlations were found between several of the inferred parameters in order to maintain the transmission potential, *R*_0_, of the Zika virus throughout the study period. A shorter inferred extrinsic incubation period was associated with reduced mosquito longevity, amplified oscillations in mosquito longevity were correlated with attenuated oscillations in mosquito density and a reduction in the minimum mosquito-to-human ratio. Pearson’s correlation coefficients, *r*, were calculated from 2,000 random samples from the posterior distribution.

### 3.3 Model forecasting

In order to highlight the model’s capacity to forecast outbreaks, the model was fit to empirical data from the 2015 epidemic only. Then, the model was simulated forward in time until the end of 2016 using parameter values inferred from the 2015 model fit and the 2016 climate data. The temporal dynamics of observed incidence from the forecast closely tracked the behaviour of the weekly notified case data (Figure 12A). However, due to the large uncertainty in forecasted incidence, there was a tendency for the number of cases to be overestimated during the 2016 outbreak. In alignment, the inferred attack rate during 2015 was on average 20% lower than estimates gained from fitting to both outbreaks (Figure 12B), indicating that in order to capture the small magnitude of the 2016 outbreak, sufficiently high levels of herd-immunity prior to 2016 was required to inhibit transmission.

**Figure 12.**
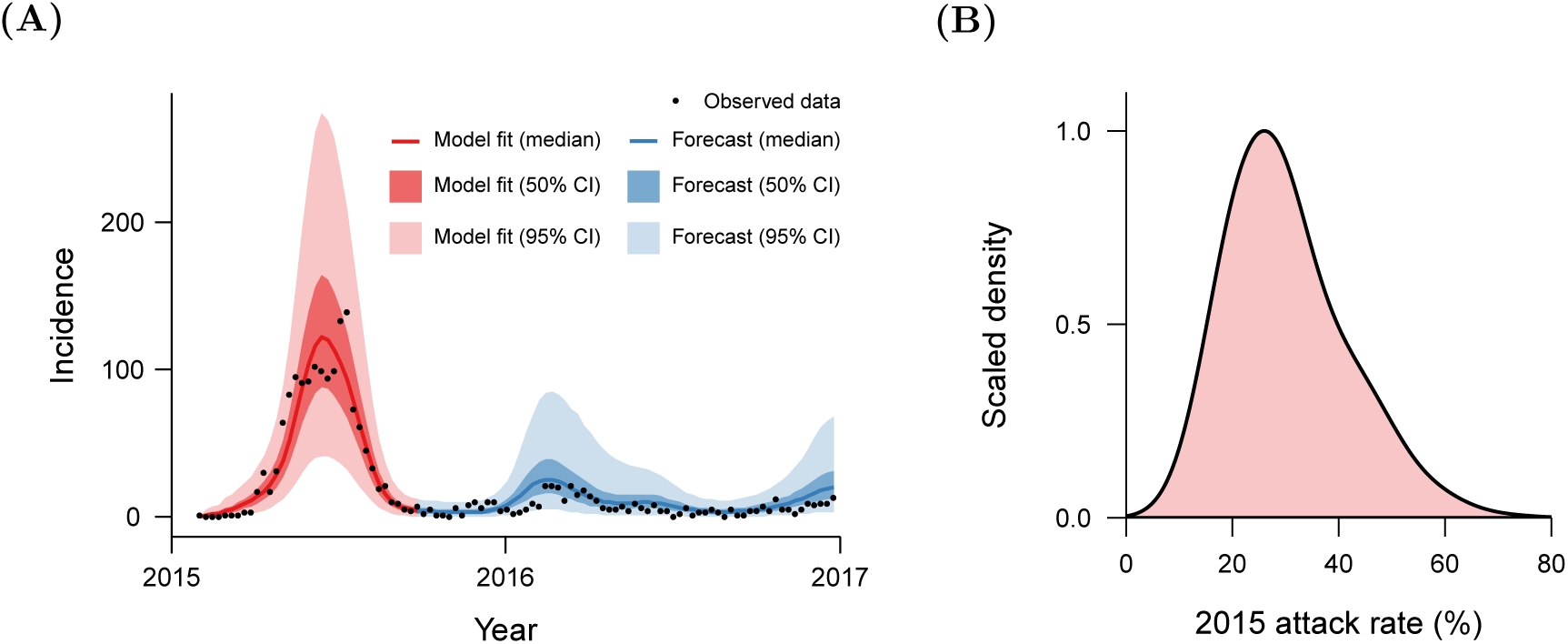
Forecasting from the 2015 epidemic. Fitting the model to the 2015 epidemic only and simulating forward in time yielded **(A)** forecasts which accurately predicted the temporal signature of the empirical data. **(B)** In turn, estimates of the percentage of the population infected during 2015 were lower than the inferred attack rate from fitting to entire empirical data set. Distributions shown were calculated from 2,000 randomly sampled values from the posterior distributions of the model fit.

### 3.4 Effects of population size on model inference

The presented model fit was executed on a human population of 100,000 individuals, whereas the full population of Feira de Santana has over 500,000 individuals (United Nations, 2015). In order to assess the influence of population size on the inference of unobserved parameters, the model was also fit with 50,000 and 500,000 individuals, keeping community size constant, and the resulting posterior distributions were compared. An increase in the number of individuals reduced the stochasticity of the simulation, and thus the variance in the estimate of the marginal distribution was reduced. Therefore, in order to optimise the pseudo-marginal method the number of particles selected at each step in the MCMC algorithm was adjusted to be *N* = 50 for 50,000 individuals and *N* = 10 for 500,000 individuals.

The majority of posterior distributions of unobserved parameters were extremely comparable between the fits for the three different population sizes (Figure 13). However, the mean and variance of the posterior distribution for the observation rate, *p*_*obs*_, was found to scale linearly with population size. This was because total infections increased with population size, thus observation rates needed to be reduced such that model generated observations matched the incidence data.

**Figure 13.**
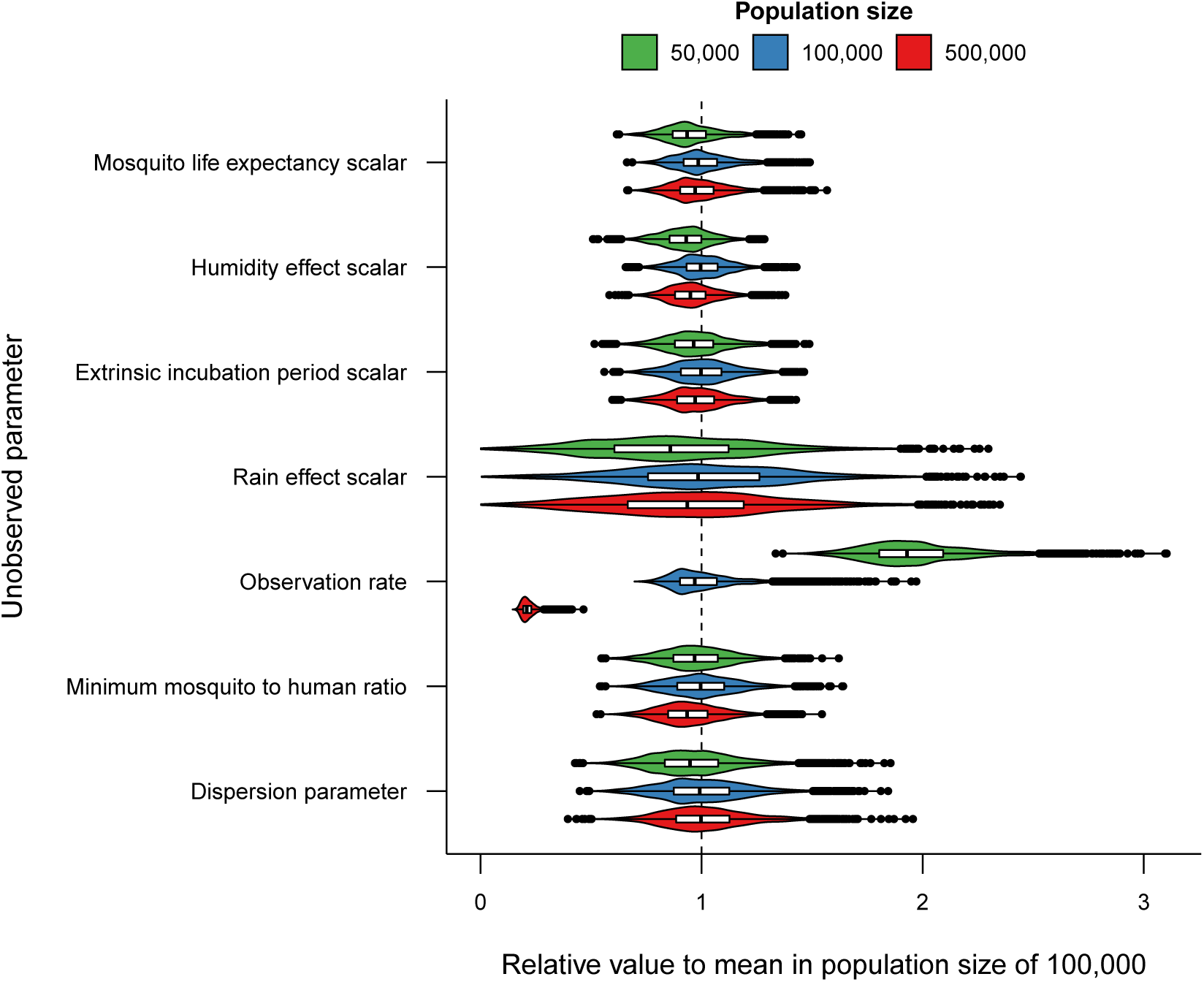
Invariance of population size on posterior distributions. The model was fit to the empirical data with three different population sizes, 50,000, 100,000 and 500,000 individuals, with fixed community size, and posterior distributions compared. The majority of posterior distributions for the three model fits were analogous with the exception of the observation rate, *p*_*obs*_, which had mean and variance that scaled by the number of individuals in each model fit.

### 3.5 Effects of mosquito mortality rates on model inference

In order to determine the effects of different assumptions about mosquito mortality rates on inferred parameters, motivated by the significant difference between *R*_0_ estimates under different assumptions of vector mortality rates (Tennant and Recker, 2018), the individual based model was fit to the empirical data under the assumption of constant vector mortality (*c*_*v*_ = 1) and then compared to the previous model fit assuming age-dependent vector mortality (*c*_*v*_ = 4).

We found that there was little difference between most posterior distributions of constant and age-dependent vector mortality. However, the inferred linear scalar that controls mean mosquito mortality rates, *η*, was higher on average under the assumption of constant daily vector mortality rates, resulting in lower mean mosquito life expectancy (Figure 14A). In order to maintain the same transmission potential, or *R*_0_, the mean infectious period of mosquitoes is required to be the same under both assumptions, which is achieved with lower mosquito life expectancies under the assumption of constant vector mortality.

**Figure 14.**
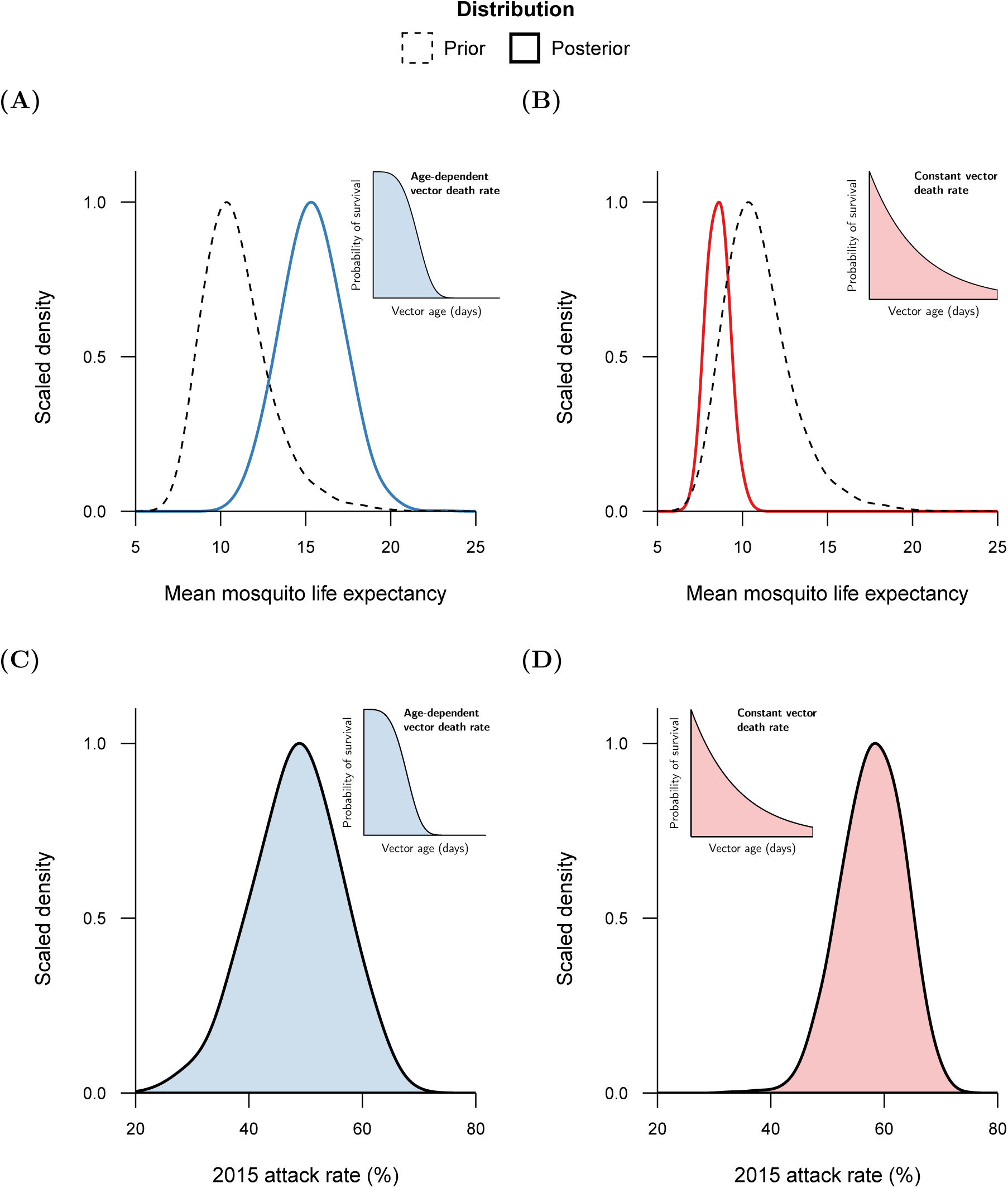
Effect of constant and age-dependent vector mortality rates on model inference. The model was fit under the assumption of constant vector mortality (*c*_*v*_ = 1) and age-dependent vector mortality (*c*_*v*_ = 4). In order to maintain transmission potential, or *R*_0_, estimates of mean mosquito longevity were reduced from the model fit with **(A)** age-dependent mortality versus the model fit with the **(B)** constant vector mortality rate assumption. Smaller variance in the posterior distribution of mean mosquito life expectancy produced decreased uncertainty in estimates of the proportion of the total population that were infected during 2015, or 2015 attack rate, from the **(C)** age-dependent fit to the model fit under the assumption of **(D)** constant vector mortality.

Interestingly, the uncertainty in the percentage of the population infected during 2015, or the 2015 attack rate, decreased under the constant vector mortality rate assumption (Figure 14B). This was due to the decrease in the variance of mean mosquito longevity, resulting in decreased variance of *R*_0_ between simulations.

### 3.6 Effects of spatial structure on model inference

To investigate the impact of space structure on model inference, the model was fit under the assumption of three lattice configurations of increasing number of communities (|*C*| = 100; 1, 000; 10, 000) with fixed local mobility and total human population size. Increasing the number of communities in the lattice, or decreasing number of individuals within each community, increased the mean minimum mosquito-to-human ratio (Figure 15A) and the effect of rainfall on mosquito density (Figure 15B). In stark contrast, the scalar controlling mean mosquito mortality rates increased as spatial resolution was refined (Figure 15C), resulting in a decrease of mean mosquito longevity throughout the study period, counterbalancing the aforementioned rise in mean mosquito density. Furthermore, there was a slight increase in the effect of humidity on mosquito life expectancy as the number of communities was increased (Figure 15D), amplifying the seasonal oscillations in mosquito longevity.

**Figure 15.**
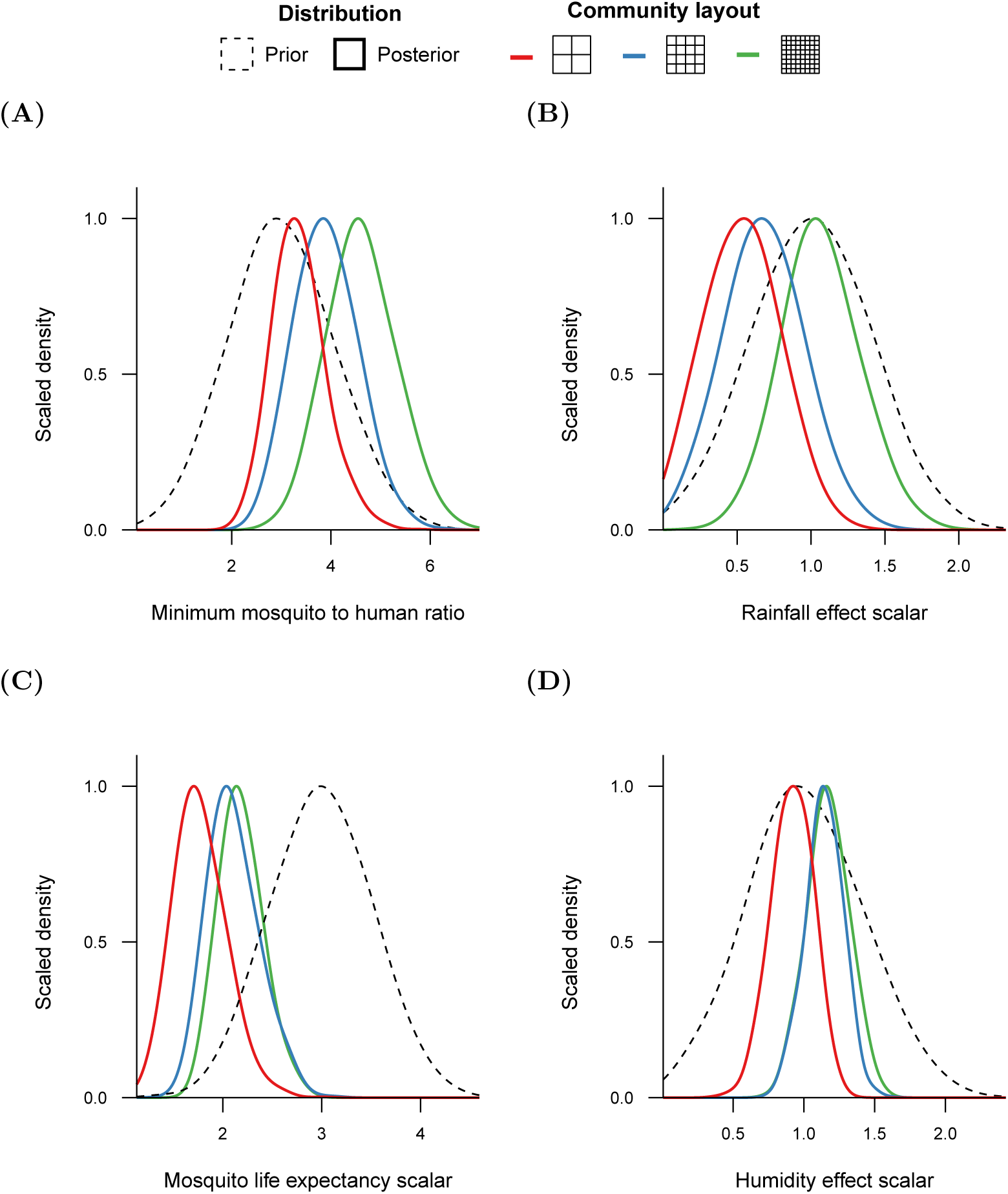
Effect of spatial refinement on model inference. The individual based model was fit to the empirical data with three different community layouts: a small lattice (|*C*| = 100), the default lattice (|*C*| = 1, 000) and a large lattice (|*C*| = 10, 000). As the size of the lattice increased, posterior distributions for **(A)** minimum mosquito density, *κ*, and **(B)** effect of rainfall on mosquito density, *ρ*_*R*_, shifted toward larger values, inferring a higher mean mosquito population density. In contrast, the scalar for **(C)** mean mortality rates, *η*, and the **(D)** effect of humidity on mortality rates, *ρ*_*H*_ increased with greater spatial refinement.

To further discern the influence of space on key drivers of transmission, the accepted unobserved parameter values were transformed into the maximum mosquito-to-human ratio, expected mosquito life expectancy, and basic reproduction number, *R*_0_, at the peak of the first epidemic, here defined as the first day of June in 2015. On average, the maximum mosquito-to-human ratio during the first epidemic increased as the number of communities in the lattice was increased (Figure 16A). At high humidity levels, the amplification of the oscillations of mosquito mortality rates from increased humidity effects counterbalanced the overall decrease in mosquito longevity as spatial dimension was refined, yielding a similar mosquito life expectancy during peak transmission (Figure 16B). Therefore, the basic reproduction number increased alongside the number of communities in the lattice (Figure 16C) because higher levels of transmission, via the increase in peak mosquito density, facilitated the virus diffusion throughout lattices of increased dimension.

**Figure 16.**
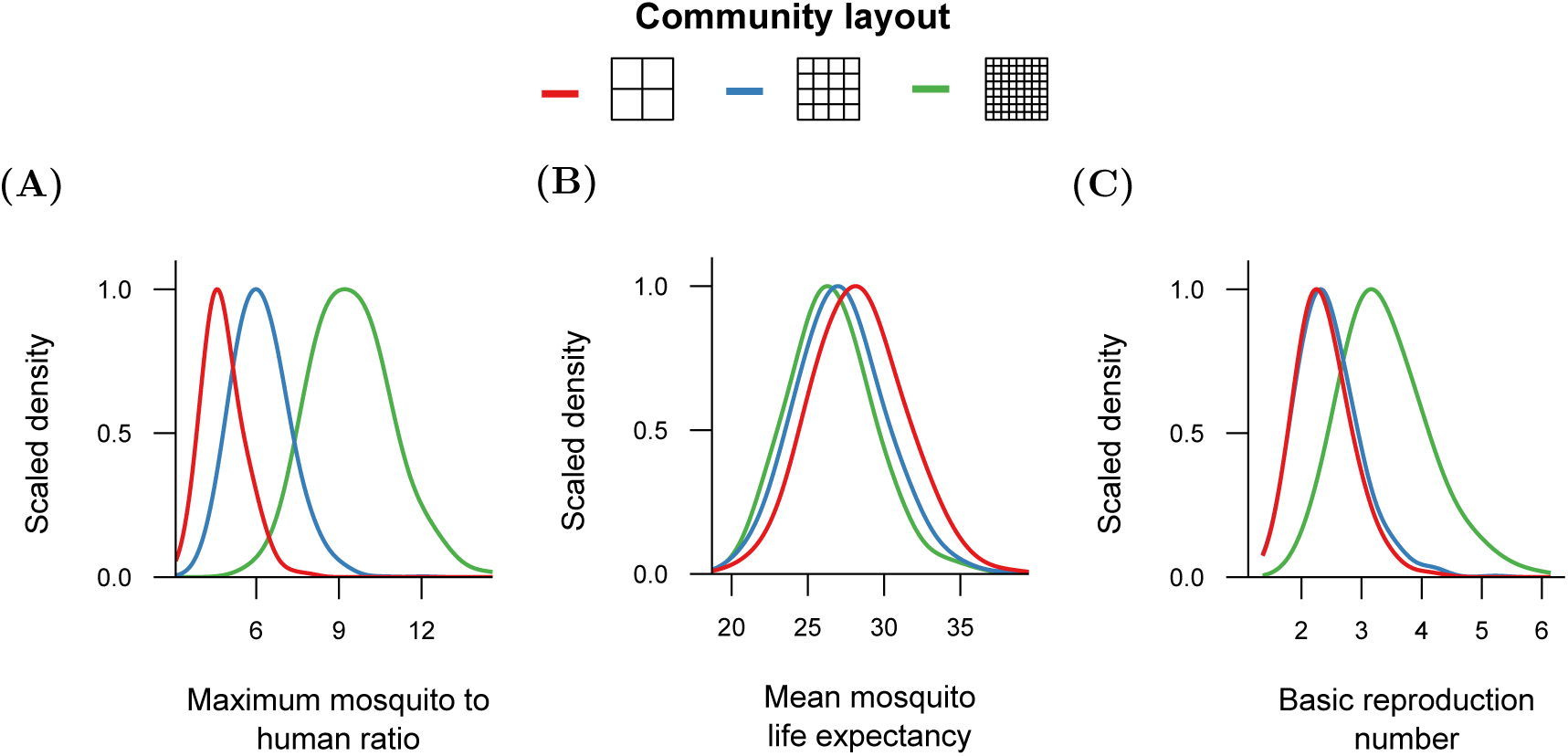
Drivers of transmission during the 2015 epidemic. Posterior distributions for estimates of key transmission parameters at peak transmission during the first epidemic, defined here as 1st June 2015, were calculated from model fits under three different community layouts: a small lattice (|*C*| = 100), the default lattice (|*C*| = 1, 000) and a large lattice (|*C*| = 10, 000). With increased spatial refinement, increased **(A)** mosquito density and consistently high **(B)** mosquito longevity yielded increased values of **(C)** the basic reproduction number, *R*_0_.

### 3.7 Effects of importation and mobility on model inference

The spatially-explicit framework further permits us to determine the influence of national and local mobility on disease outbreaks, so the model was fit to the empirical data with three fixed values for external introduction rates (*ι* = 0.5, 1, 5) and three levels of local mobility (1 − *p*_*σ*_ = 0.25, 0.5, 0.75).

All model fits exhibited shifts in the posterior distribution of each unobserved parameter. Critically, there was a bistable response in inferred seroprevalence levels at the end of the two year study period. High external introduction rates, in combination with any level of local mobility, consistently produced immunity levels of over 50% (Table 6). This was in stark contrast to inferred immunity levels of less than 10% for lower introduction rates and local human mobility. However, high local mobility also consistently inferred human seroprevalence of over 50%, as the virus could disseminate throughout the community structure more easily.

**Table 6.** Bi-stability of inferred seroprevalence. The individual based model was fit under three external introduction rates, *ι* = 0.5, 1, 5, and three levels of local mobility, 1 − *p*_*σ*_ = 0.25, 0.5, 0.75. All model fits with high external infection rates or high local mobility were consistently attracted towards posterior distributions of high immunity levels of greater than 50%. However, fits lower introduction rates and local mobility hindered transmission and inferred very low attack rates of less than 10% from February 2015 to December 2016. Results shown are the median and 95% credible intervals of human seroprevalence at the end of 2016.

|  |  | Local mobility |  |  |
| --- | --- | --- | --- | --- |
|  |  | Low | Medium | High |
| External infection rate | High | 58.8%<br>[41.4%, 69.7%] | 59.2%<br>[44.9%, 68.5%] | 61.1%<br>[52.4%, 68.7%] |
|  | Medium | 6.0%<br>[4.4%, 8.1%] | 6.2%<br>[4.5%, 8.4%] | 59.6%<br>[51.6%, 67.4%] |
|  | Low | 3.1%<br>[2.2%, 4.2%] | 3.0%<br>[2.1%, 4.3%] | 58.8%<br>[50.1%, 65.4%] |

Model fits inferring high seroprevalence produced high values of *R*_0_ prior to the 2015 outbreak before falling below one in July 2015 (Figure 17A). In contrast, inferred *R*_0_ estimates were much smaller in model fits that estimated low seroprevalence (Figure 17B). There was little difference between *R*_0_ estimates at peak levels of transmission across all tested combinations of external introduction rate and local mobility, however, which implies that lower seroprevalence estimates were induced simply by shorter durations of high transmission potential.

**Figure 17.**
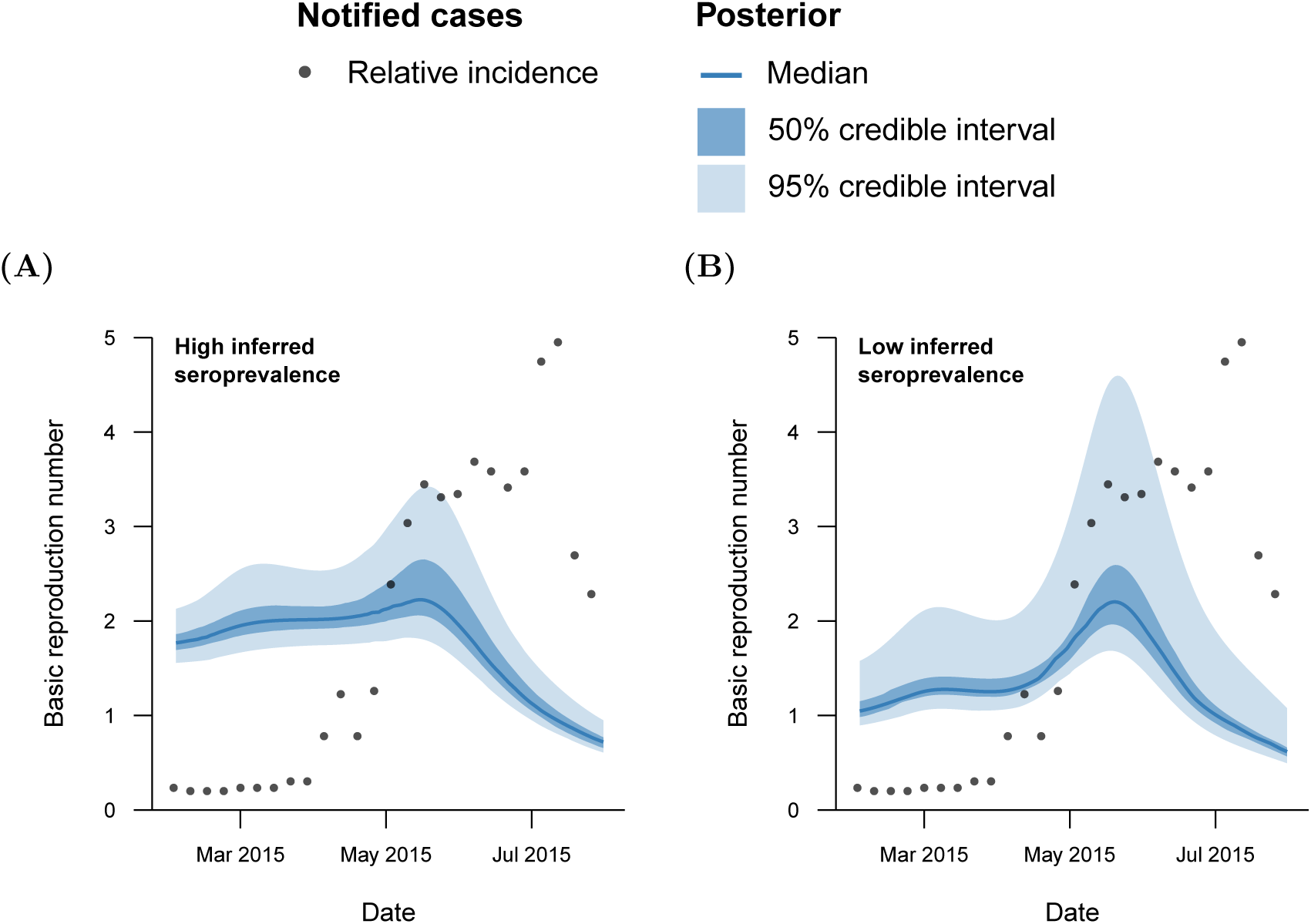
Effect of inferred seroprevalence on the basic reproduction number R_0_. **(A)** High external introduction rates or local mobility (here, *ι* = 1 and 1 − *p*_*σ*_ = 0.75) consistently inferred high seroprevalence due to sustained levels of *R*_0_ from February 2015 until July 2015. **(B)** In contrast, model fits inferring low immunity levels (here, *ι* = 1 and 1 − *p*_*σ*_ = 0.5) exhibited *R*_0_ estimates which were almost singular prior to May 2015, hindering transmission. Median and credible intervals shown were calculated from 1,000 randomly sampled values from the posterior distribution of each model fit.

Spatially, high external introduction rates (*ι* = 5) and low local mobility (1−*p*_*σ*_ = 0.25) generated hundreds of outbreaks which slowly spread locally over the first epidemic in 2015. A large number of small clusters of high susceptibility provided ideal conditions for virus invasion during the second smaller outbreak, yielding a heterogeneous immunity landscape (Figure 18A). As the external introduction rate was lowered (*ι* = 0.5), the number of initial cases in 2015 greatly decreased. Hindered by low local mobility, very high susceptibility levels were maintained throughout the lattice that failed to be penetrated during the 2016 outbreak due to poor external introduction and local mobility (Figure 18B). However, increased local mobility (1 − *p*_*σ*_ = 0.75) enabled the small number of initial cases to rapidly spread throughout the lattice, creating large spatial clusters of high immunity and susceptibility. During the second peak, neighbourhoods of complete susceptibility permitted further local expansion of the virus, creating an overall landscape of moderate immunity with collections of entirely immunised communities (Figure 18C).

**Figure 18.**
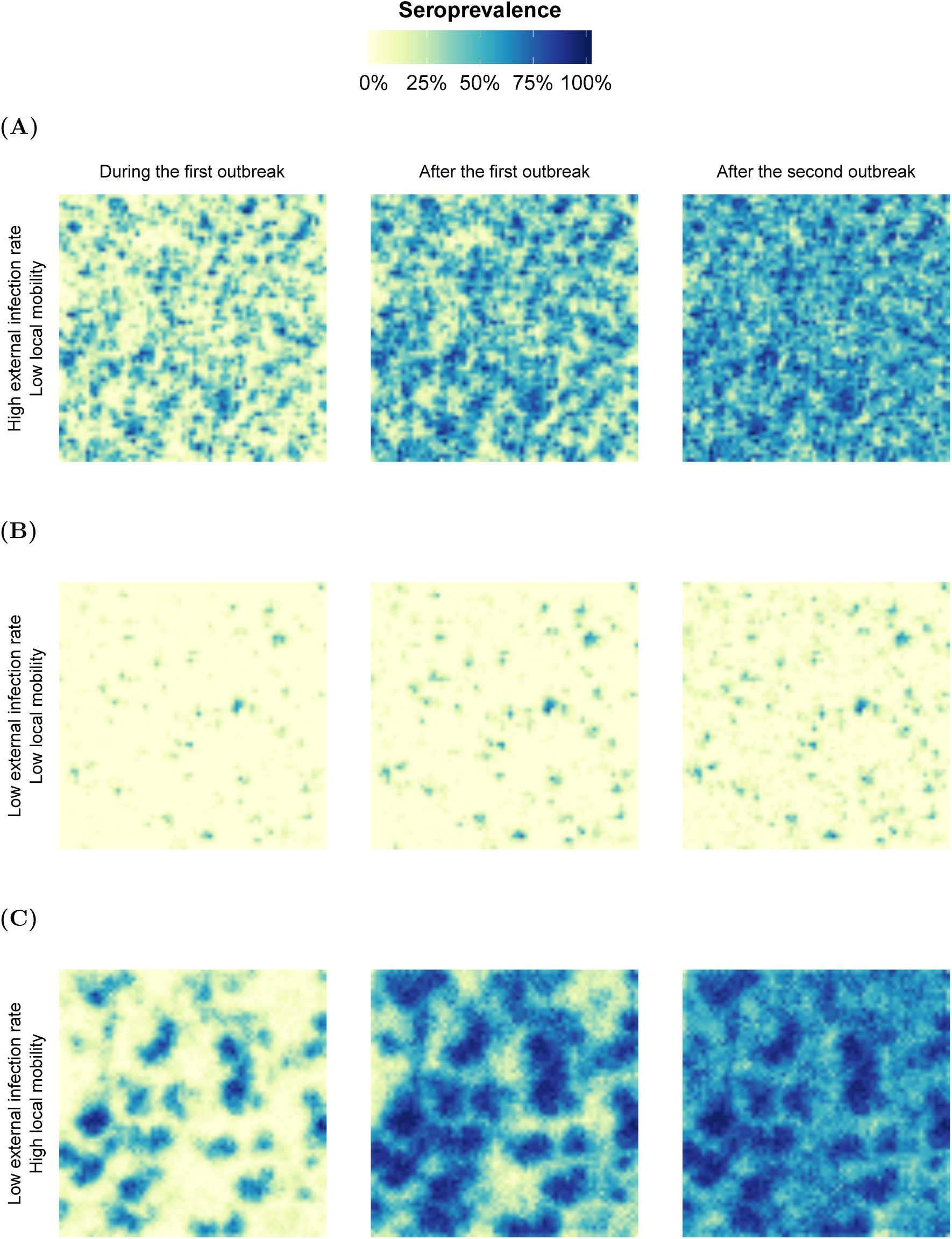
Effect of mobility and external introduction rates on Zika spatial dynamics. The individual based model was fit under different assumptions of local and global mobility and accumulation of immunity across the lattice was observed. **(A)** High external introduction rates (*ι* = 5) and poor human mobility (1 − *p*_*σ*_ = 0.25) gave rise to many initial outbreaks culminating in a highly heterogeneous landscape of immunity. **(B)** Low local mobility in combination with poor introduction rates (*ι* = 0.5), greatly hindered spatial spread of the virus, yet **(C)** increasing human mobility (1 − *p*_*σ*_ = 0.75) enabled rapid diffusion of the virus throughout the lattice. Seroprevalence was calculated from the percentage of individuals within each community with acquired immunity to the virus.

## 4 Discussion

Climate has previously been shown to play a crucial role in the emergence and spread of arboviral disease (Huber et al., 2018; Lowe et al., 2013, 2016; Tesla et al., 2018). However, the exact relationships between climate and epidemiological drivers are not yet well established in the field. Therefore, here, we sought to quantify the influence of climate on vector suitability and virus transmissibility by fitting a spatially-explicit, climate-dependent individual based model to incidence data. A computational speed-up in the individual based model allowed us to perform model fitting within a fully Bayesian framework for the first time.

The model was first fit to simulated data from the individual based model itself. Fitting to the simulated data showed that key epidemiological transmission parameters, such as mosquito longevity, could be reliably inferred given that all other parameters were well-informed. We also found correlations between several inferred parameters, such as the effects of rainfall and humidity on mosquito density and longevity. This was confounded by the strong relationship between rainfall and humidity within the climate data, suggesting potential for model simplification.

These findings were consistent with our model fit to Zika incidence data from Feira de Santana, Brazil. For the empirical data, the framework identified posterior distributions of parameters, defining relationships between climate and mosquito demography. These results generally indicate a strong influence of humidity on mosquito longevity and a relatively weaker effect of rainfall on mosquito density. However, we found that there was little impact of annual oscillations in temperature on the Zika outbreak within this region. This was likely due to the fact that temperatures were high throughout the study period, thereby enabling short extrinsic incubation periods of the virus.

Crucially, these findings aligned with a climate-driven ordinary differential equation model fit by Lourenço et al. (2017), including inferred reporting rates of less than 1%. These reporting rates were in stark contrast to estimated Zika reporting rates of 16% and 18% in El Salvador and Suriname, respectively (Shutt et al., 2017), but in agreement with observation rates of 2.7% in Cabo Verde Islands, West Africa (Lourenço et al., 2018). High seroprevalence levels of over 50% were also agreed with previous cross-sectional serological studies in Salvador, Brazil (Netto et al., 2017), Yap Island (Duffy et al., 2009), French Polynesia (Cauchemez et al., 2016), and Nicaragua (Zambrana et al., 2018). This combination of low observation and high attack rates suggest great potential for asymptomatic infected individuals to transmit the virus. Indeed, Zika, as has dengue, has been shown to have a high proportion of asymptomatic cases (Haby et al., 2018; Ladhani et al., 2016), although it is not yet clear whether asymptomatic and symptomatic individuals have the same transmission potential (Duong et al., 2015; Moghadas et al., 2017; ten Bosch et al., 2018).

Traditional modelling frameworks, such as the one by Lourenço et al. (2017), implicitly assume constant vector mortality rates, which have been shown to impact the basic reproduction number, *R*_0_, of arboviral diseases (Tennant and Recker, 2018). Due to the flexibility of the individual based model, the model was fit to the empirical data under constant and age-dependent mosquito mortality rates. Under both assumptions, mean mosquito life expectancies were within previously found bounds of one to two weeks (Maciel-de Freitas et al., 2007; Marinho et al., 2016; Muir and Kay, 1998). However, in order to maintain the same basic reproduction number, the inferred mosquito life expectancy greatly shortens under the assumption of constant death rates.

The importance of model assumptions for inferring parameter values was also highlighted by the fact that community structures of finer spatial resolution required increased effects of rainfall on mosquito density to capture the explosive dynamics of the 2015 outbreak. A similar behaviour was also found when fitting the model under different local human mobility and importation rates. Dependent on these rates, inferred seroprevalence levels at the end of 2016 exhibited a bistable behaviour. For high seroprevalence, external introduction rates were required to be high. This implies that the introduction of infected individuals into multiple locations within Feira de Santana was crucial in the emergence of Zika during the 2015 epidemic. From a public health perspective, identifying common socio-ecological features in these locations could therefore be useful in determining infection risk factors and informing outbreak prevention strategies.

To that end, we further investigated the potential of the model to be used in disease forecasting by fitting exclusively to the 2015 epidemic and simulating forward in time. Our predictions matched the temporal signature of the 2016 empirical data, and demon-strated that reliable climate data would be required in order to accurately predict disease outcomes. That is, although climate factors exhibit general annual trends, these alone are likely insufficient to forecast incidence accurately. Due to the high variance of inferred attack rates, the framework would also benefit from cross-sectional serological data in order to more precisely quantify the relationships between climate and vector suitability. This may not be enough in cases where climate drivers are strongly correlated, such as here. In these cases, more robust estimates of some model parameters, such as mosquito longevity, could go a long way in more accurately quantifying ecological features that are challenging to measure empirically, such as the mosquito carrying capacity.

With that in mind, many epidemiological and demographical parameters were fixed within our framework. Inference on the parameters of interest was therefore dependent upon the choices of these fixed parameters. For example, we demonstrated the impact of different fixed human mobilities on the spatio-temporal dynamics of disease incidence. This suggests the strong benefits that including spatio-temporal incidence or social data sets could have on assessing the importance of human movement on arboviral disease outbreaks. It should also be noted that the model is still relatively computationally expensive in comparison to a spatially homogeneous deterministic system. Model fitting run-times could further be reduced by instead using Particle marginal Metropolis Hastings methods to estimate the posterior distribution (Andrieu et al., 2010). However, with the alleviation of computational costs and invariance of inferred climate drivers to the number of individuals within the model, it may already be practical to use an individual based model as a real-time disease control management tool, at least on time scales longer than a week.

Here, we have clearly demonstrated that not only is it possible to fit an individual based model to relatively sparse data within a fully Bayesian framework, but we can quantify relationships between climate and epidemiological features, such as mosquito longevity or the extrinsic incubation period. Unlike previous modelling approaches, the spatially-explicit nature of our framework showed that the virus was likely introduced into multiple spatial foci in order to create the observed temporal dynamics and rapid accumulation of Zika immunity in Feira de Santana. These findings emphasise the added benefits that cross-sectional serological and spatio-temporal incidence data sets could bring in more precisely inferring the ecological drivers of arboviral epidemiology. Overall, our results strongly indicate the significant impacts that spatio-temporal ecological heterogeneities have on mosquito-borne disease inference, and should thus be explicitly considered when informing control efforts through mosquito elimination and vaccine deployment programs.

